# Bacterial histone-like proteins released during antibiotic treatment mediate vascular injury in meningococcal sepsis

**DOI:** 10.64898/2026.07.15.738198

**Authors:** Dagmara McGuinness, Carla Johnson, Leandro Lemgruber Soares, Fadil Bidmos, Edwin A Yates, Jeremy Turnbull, Thomas Evans, Rebecca McHugh, Andrew J Roe, Gillian Douce, Andrew Smith, Roisin Ure, Cheng-Hock Toh, Simon Abrams, Jethro Herberg, Victoria J Wright, Ruud Nijman, Robert S Heyderman, Saul N. Faust, Michael Levin, Christopher A. Moxon

**Affiliations:** School of Infection and Immunity, MVLS, University of Glasgow; Glasgow, G12 8TA, UK; Section of Paediatrics, Department of Infectious Diseases, Faculty of Medicine, Imperial College London; London, SW7 2AZ, UK; Department of Biochemistry, Institute of Systems, Molecular and Integrative Biology, University of Liverpool; Liverpool, L69 7ZX, UK; Intellihep, Liverpool L69 7ZB, UK; Copenhagen Centre for Glycomics, Institute of Cellular and Molecular Medicine, University of Copenhagen; Blegdamsvej 3, 2200 København N, Copenhagen, Denmark; School of Medicine, Dentistry & Nursing, MVLS, University of Glasgow; NHS GGC SmiRL, Glasgow, G12 8QQ, UK; Clinical Infection, Microbiology & Immunology, University of Liverpool; Liverpool, L69 3BX, UK; Research Department of Infection, Division of Infection and Immunity, University College London; London, WC1E 6BT, UK; NIHR Southampton Clinical Research Facility and Biomedical Research Centre, University Hospital Southampton NHS Foundation Trust, and Faculty of Medicine and Institute for Life Sciences, University of Southampton; Southampton, SO16 6YD, UK

**Author notes:** Contact during submission and.

**Keywords:** *Neisseria meningitidis*, meningococcal sepsis, histone-like DNA-binding proteins, endothelial damage, non-anticoagulant heparins, *Galleria mellonella*

## Abstract

Vascular injury and coagulopathy are key drivers of mortality in bacterial sepsis. In *Neisseria meningitidis* infection, endothelial adhesion and thrombosis cause the characteristic petechial rash and, in the most severe cases, purpura fulminans. Although antibiotics rapidly kill bacteria, inflammation and vascular injury often persist or worsen after bacterial clearance, suggesting ongoing toxicity from released bacterial components. Here we identify bacterial histone-like proteins (HLPs), small positively charged DNA-binding proteins conserved across bacterial species, as previously unrecognized mediators of vascular damage. *In vitro* HLPs are released following antibiotic exposure, disrupting endothelial integrity. In patients with severe sepsis, they are detectable in plasma and tissue, colocalising with areas of vascular leak and coagulopathy. Non-anticoagulant heparins and anti-HLP antibodies neutralize HLP-induced endothelial disruption and toxicity *in vitro* and *in vivo*. These findings reveal HLPs as antibiotic-released bacterial toxins and suggest new therapeutic strategies to prevent vascular injury in sepsis.

**One sentence summary:** Antibiotic treatment of meningococcal and other sepsis causative bacteria triggers the release of histone-like proteins (HLPs) that damage the endothelium and drive coagulopathy, but these effects can be neutralized by non-anticoagulant heparins or anti-HLP antibodies.

## Introduction

Vascular damage and coagulopathy are hallmarks of bacteraemic sepsis, which contributes to approximately 11 million sepsis-related deaths annually worldwide (*1*). In *Neisseria meningitidis* infection, thrombosis and vascular leakage cause the characteristic petechial rash (*2*). In purpura fulminans, the most severe form, widespread lesions lead to ischemia, necrosis, limb loss, and high mortality (*3*). These outcomes reflect both a high bacterial load and the ability of *N. meningitidis* to adhere to and dysregulate vascular endothelial cells (*2, 4, 5*). Type IV pili mediate endothelial adhesion and contribute endothelial damage and coagulation activation, by inducing shedding of the anti-coagulant protein Endothelial Protein C Receptor (EPCR). However, while potentially druggable (*6–8*) anti-pili drugs are yet to be tested in septic patients.

Lipopolysaccharide (LPS) endotoxin is another major virulence factor that triggers immune activation and cytotoxic responses across multiple cell types (*9*). Despite extensive development of endotoxin-neutralizing therapies, including those reaching clinical trials, none have improved survival or entered clinical use (*10–12*). Host-focused therapies have also been tested without any yet being adopted in clinical practice guidelines. Activated Protein C, once thought promising (*13*), was withdrawn after a multicentre study showed increased bleeding risk and no mortality benefit (*14*).

These limitations highlight the need for alternative therapeutic approaches. Although antibiotics rapidly eliminate *N. meningitidis* and other sepsis-causative bacteria, inflammation and clinical deterioration often persist or worsen during and after bacterial clearance. This seeming paradox, observed in both patients and experimental models, likely reflects release of proinflammatory and cytotoxic bacterial components upon bacterial killing (*15*). In a murine sepsis model, antibiotics failed to reduce mortality, and similar clinical outcomes occurred following transfer of antibiotic-killed bacteria (*16*). These findings reveal a temporal therapeutic window for adjunctive treatments that neutralize bacterial products released during antibiotic therapy. While LPS endotoxin has been the primary focus (*17–19*), its neutralization alone has not proven sufficient, suggesting the involvement of additional bacterial factors contributing to endothelial injury and coagulopathy warrant investigation.

We hypothesized that bacterial histone-like proteins (HLPs); small and positively charged, nucleoid-associated DNA-binding proteins conserved across bacteria, may represent such factors. This concept parallels findings where extracellular histones, released from dying or activated cells, are potent mediators of endothelial damage, coagulation activation, and vascular leakage through charge-dependent membrane interactions (*20–22*). Host released extracellular histones are established mediators of sepsis-associated vascular injury (*23, 24*). Analogously, in *Plasmodium falciparum* malaria, parasite-derived histones induce endothelial damage and coagulopathy at sites of parasite sequestration (*25*), where adherent histones were visualized on the luminal surface of the endothelium, suggesting that pathogen accumulation can concentrate the release of histones to the endothelium (*26*). Although bacterial HLPs differ in sequence from mammalian histones, their strong positive charge and small size suggest similar pathogenic potential.

HLPs are known to be surface-exposed and secreted by several bacterial species, including *Streptococcus pyogenes* and Gram-negative pathogens, where they bind to negatively charged components such as LPS and DNA within biofilms (*27, 28*). Collectively, these findings suggest that HLPs may be released into the host environment during infection or bacterial lysis. HLPs have also been implicated in inflammatory disease: anti-HLP antibodies are detected in patients with primary biliary cirrhosis, associated with chronic inflammatory liver lesions (*29*). In vitro, *S. pyogenes* HLPs activate monocyte inflammatory signalling (*30, 31*), further supporting their role as proinflammatory mediators. However, it remains unknown whether HLPs are released following antibiotic treatment or detectable in patients with meningococcal or other forms of sepsis. Their direct effects on endothelium and contribution to vascular pathology have also not been defined.

Here, using a combination of *in vitro* and *in vivo* approaches and patient samples we establish that HLPs are released after antibiotic treatment and cause endothelial barrier disruption and are present in patients with meningococcal sepsis, co-localising with areas of leak and coagulopathy. In both *in vitro* and *in vivo* model, non-anticoagulant heparins and anti-HLP polyclonal antibodies neutralise HLP-induced endothelial disruption and toxicity. These findings identify HLPs as previously unrecognized bacterial mediators of vascular injury and suggest new therapeutic strategies for limiting endothelial damage and coagulopathy in sepsis.

## RESULTS

### HLPs released from meningococci by antibiotic treatment disrupt endothelial barrier

First, we aimed to determine whether meningococcal HLPs are released into bacterial culture medium and whether this is enhanced by antibiotic treatment. We used the *N. meningitidis* serogroup B wild-type strain H44/76 (subsequently referred to as wild-type) and treated it with ceftriaxone, a third-generation cephalosporin widely used as first-line treatment for paediatric sepsis. Using affinity-purified polyclonal antibodies against *N. meningitidis* serogroup B (MC58) HLPs [DNA-binding protein HU-beta (hupB), Integration host factor subunit beta (IHFβ) and Integration host factor subunit alpha (IHFα)], we detected low levels of HLPs in the culture supernatants of untreated meningococci. Upon ceftriaxone treatment, we observed a dose-dependent release of hupB and IHFβ into the culture medium **(Fig. 1A, fig. S1A)**.

**Figure 1.**
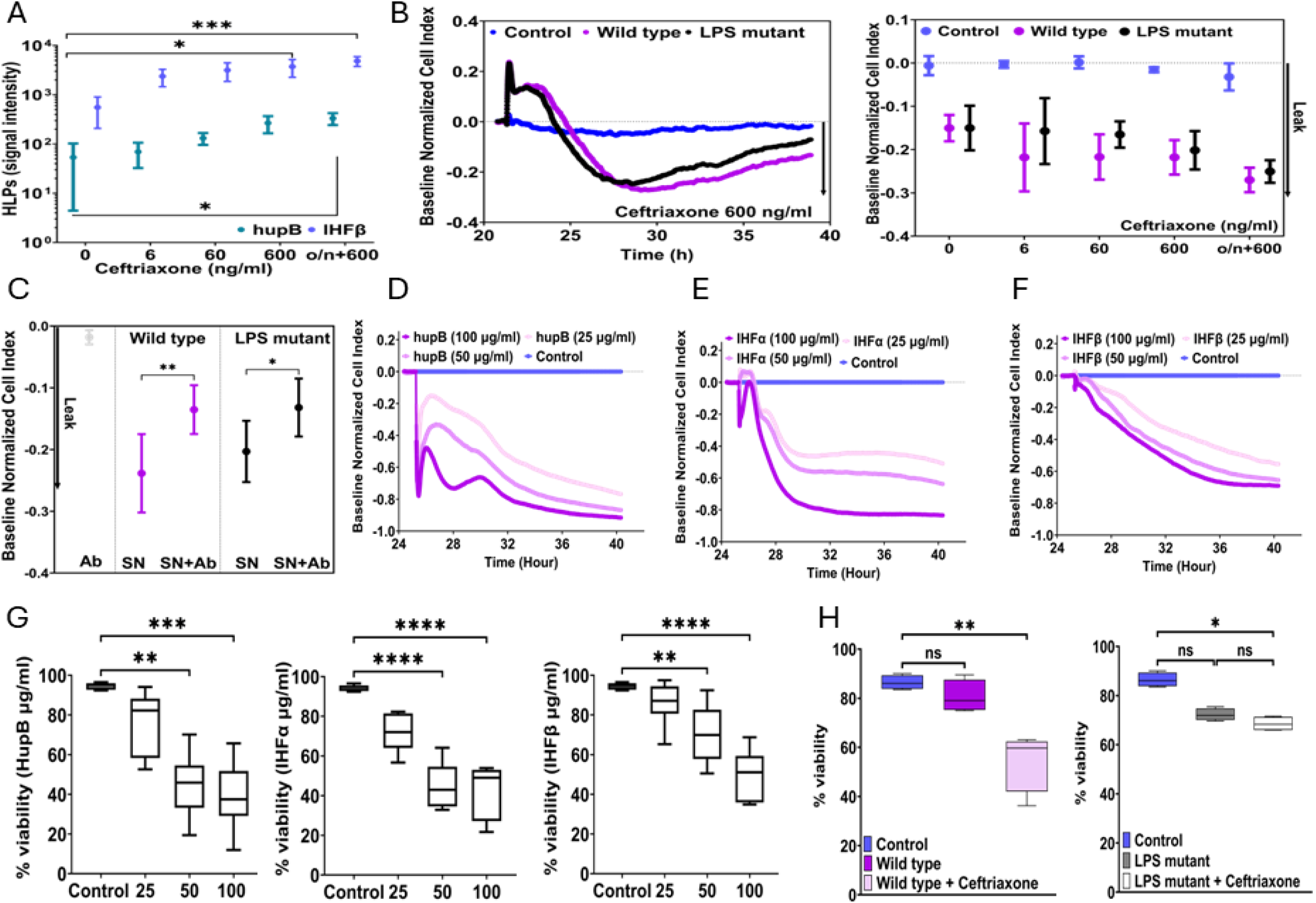
HLPs released by antibiotic treatment and *in vitro* are necessary and sufficient to cause endothelial barrier disruption. **(A)** Dose-dependent release of HLPs into the culture supernatant after Ceftriaxone treatment was analysed using Western blot and custom anti-HLPs antibodies, **(B)** The supernatants from *Neisseria meningitidis*, strains H44/76 (wild type) and H44/76[pHBK30] (LPS-mutant) induce endothelial monolayer leak in a dose-dependent manner. Cellular indexes are normalised to vehicle. Left chart: An example of dermal endothelial cell traces in the response to *N. meningitidis* strains treated with 10x minimum inhibitory concentration (MIC) of Ceftriaxone. Right chart: endothelial monolayer permeability (leak) is dose-dependent, **(C)** Disruptive effect of bacterial supernatants from both meningococci strains is abrogated by pre-incubation with anti-HLPs antibodies before adding to endothelial cells. Recombinant HLPs: **(D)** hupB, **(E)** IHFα and **(F)** IHFβ induce endothelial monolayer leak in the dose-dependent manner, (**G)** Recombinant HLPs influence endothelial cells viability in dose-dependent manner, (**H)** *Neisseria meningitidi*s supernatants from both strains affect endothelial cell viability. Experiments were performed in triplicate. P<0.005 was considered significant. *P< 0.05, **P < 0.01; ****P< 0.0001; ns non-significant. Data presented as mean with SD.

Next, we investigated whether HLPs released from antibiotic-exposed meningococci could disrupt endothelial barrier function using an impedance-based system (xCELLigence). Meningococcal supernatant disrupted the barrier function of primary dermal endothelial cells (HDECs) within three hours, plateauing at six hours **(Fig. 1B)**. This effect intensified with increasing antibiotic doses applied to the meningococci. Ceftriaxone alone had no impact on endothelial integrity, even at high concentrations **(Fig. 1B)**.

To assess the contribution of endotoxin to barrier disruption, we used an H44/76 LPS mutant [pHBK30], which is viable but lacks detectable LPS and has no endotoxin activity (*32*). At the same bacterial and antibiotic concentration, the LPS-mutant caused the same degree of barrier disruption as the wild-type strain **(Fig. 1B)**. Similarly, whole meningococci killed with supra-MIC doses (6 ng/ml) of ceftriaxone exhibited potent and comparable barrier-disruptive effects, regardless of endotoxin activity or antibiotic concentration **(fig. S1B, S2)**. The absence of endotoxin in the LPS-mutant strain was previously well-characterized (*32*) and further confirmed by sequencing and by detecting endotoxin in the wild-type but not in the LPS- mutant **(fig. S3)**. In contrast, activation of endothelial cells, measured by the induction of the intracellular adhesion molecule (ICAM-1), was significantly higher following exposure to supernatants or whole bacteria from wild type meningococci compared to the LPS-mutant strain **(fig. S4)**.

Together these data indicate that, under these *in vitro* conditions in endothelial cells, LPS endotoxin is necessary for maximal inflammatory effect but is not for the barrier-disruptive effects of meningococcal supernatants or whole antibiotic-killed meningococci. To determine whether HLPs alone are barrier disruptive, we removed hupB, IHFα, and IHFβ from the supernatants prior to the treatment via immunoprecipitation using beads coated with a cocktail of affinity-purified polyclonal antibodies. The resulting HLP-depleted supernatants exhibited significantly reduced barrier-disruptive effects **(Fig. 1C)**, suggesting that HLPs, either alone or in protein complexes, contribute to endothelial barrier dysfunction. Treatment of the supernatants with DNase I did not mitigate this effect, indicating that bound DNA was not responsible for the disruption **(fig. S5)**. To assess whether HLPs were sufficient to cause barrier disruption and to compare their effects to eukaryotic extracellular histones, we generated recombinant HLPs (rHLPs) for hupB, IHFα, and IHFβ. Each rHLP increased endothelial barrier permeability in a dose-dependent manner at concentrations above 25 μg/ml **(Fig. 1D- F)**. LPS levels and DNase I treatment of the recombinant HLPs had no effect on barrier integrity **(fig. S5A,B)**, indicating against LPS or DNA contamination being responsible for the disruption. Supernatants from antibiotic treated bacteria as well as recombinant HLPs (at concentrations ≥50 μg/ml) but not at lower concentrations **(Fig. 1G, H)** significantly reduced cell viability, suggesting that mechanisms other than cell death likely contribute to barrier disruption at lower concentrations.

### HLPs disrupt endothelial junctions and anti-coagulant function by receptor shedding

Endothelial junctions are anchored by vascular endothelial cadherin (VE-cadherin), and its redistribution has been shown to be critical in vascular leak induced by various stimuli. To assess the potential role of VE-cadherin in barrier disruption by HLPs, we examined its distribution in endothelial cells exposed to supernatants from antibiotic-treated meningococci or recombinant HLPs. In untreated cells, VE-cadherin formed a continuous junctional network between adjacent cells. However, in cells treated with bacterial supernatants from either the wild type or LPS-mutant strain, as well as those treated with rHLPs, VE-cadherin displayed a patchy distribution with significant loss of junctional staining (**Fig. 2A, Movies S1-S3)**. LPS endotoxin was not the primary driver of junctional loss, as junctional integrity was similarly disrupted in cells treated with the LPS-mutant lacking endotoxin activity. Quantitatively, the proportion of VE-cadherin positive cells was reduced following treatment with all three rHLPs as compared to control **(Fig. 2B)**. Another hallmark of endothelial dysfunction in meningococcal disease is a shift toward a procoagulant phenotype, characterized by reduced expression of receptors involved in the anticoagulant endothelial protein C pathway (*2*). To investigate whether HLPs contribute to this effect, we assessed the distribution and expression of endothelial protein C receptor (EPCR). Like VE-cadherin, EPCR exhibited marked redistribution, with both qualitative **(Fig. 2A)** and quantitative loss of surface expression in cells treated with recombinant HLPs **(Fig. 2B)**.

**Figure 2.**
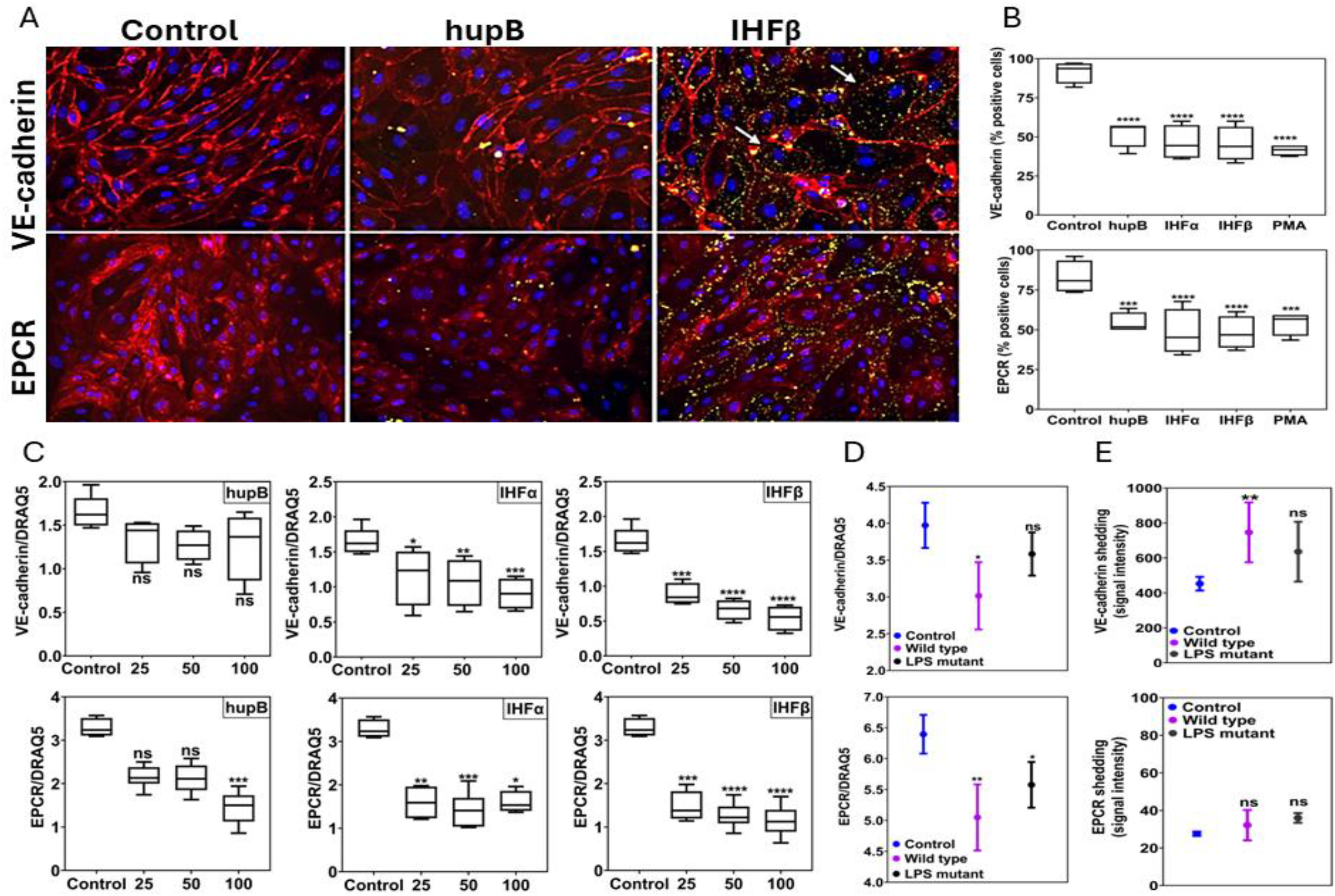
HLPs disrupt endothelial junctions and coagulation function by inducing receptor shedding. **(A)** Confluent HDECs were treated with 25ug/ml of the different HLPs for 6h and stained with fluorescently labelled antibodies raised against VE-cadherin and EPCR (see Methods). White arrows indicate leak. Magnification 20x, **(B)** The percentage of VE-cadherin and EPCR positive cells was calculated after treatment with the recombinant HLPs. IF images were scored using QuPath software, **(C)** Reduction of VE-cadherin and EPCR in response to recombinant HPLs treatment was confirmed using in cell western. HDECs were treated with 2x serial dilutions of each HLP (range 100 – 0.8 ug/ml) for 6h. Surface markers staining was normalised to the DRAQ5 staining, reflecting cell number, **(D)** Reduction of VE-cadherin and EPCR levels on the human dermal endothelial cells, in response to both bacterial strains supernatants after treatment with 10xMIC of Ceftriaxone was observed. Simultaneously, increased levels of VE-cadherin and EPCR in the endothelial cell media was noted, however this increase was not significant for EPCR (E). p<0.05 was considered as significant. *P < 0.05, **P < 0.01; ****P < 0.0001; ns non-significant. Data were presented as mean with SD.

Reduced surface VE-cadherin and EPCR could reflect internalisation, decreased production or shedding. To further examine these mechanisms, we quantified total cellular protein levels using immunofluorescence (In-Cell Western) and normalized the values to the total number of cells per well **(Fig. 2C)**. Additionally, we assessed shedding by measuring soluble protein levels in endothelial culture supernatants via Western blot. Total cellular levels of VE-cadherin and EPCR were significantly reduced following treatment with IHFα and IHFβ, even at 25 μg/ml. hupB treatment reduced surface EPCR levels at 100 μg/ml (p<0.05) but did not significantly affect surface VE-cadherin **(Fig. 2C)**. Treatment with meningococcal supernatants also decreased total cellular levels of these proteins **(Fig. 2D)**. Shed soluble VE-cadherin was higher following application of supernatant from wild type with a non-significant trend for LPS mutant meningococcal culture supernatant. Shedding of VE-cadherin was increased following treatment with supernatants from wild type meningococci, with a non-significant trend observed for the LPS mutant strain **(Fig. 2E)**. Additionally, application of IHFβ increased soluble VE-cadherin levels, whereas hupB and IHFα had no such effect. Soluble EPCR levels in endothelial culture supernatants were not significantly altered by treatment with meningococcal supernatants (wild type or LPS mutant) or any of the three HLPs **(Fig. 2E)**. These findings suggest that HLPs reduce endothelial levels of key junctional and anticoagulant proteins, likely by receptor shedding at least for VE-cadherin.

### HLPs are enriched in meningococcal skin lesions leak areas

To assess the potential clinical relevance of a role of HLPs in meningococcal disease, we measured their levels in the plasma of patients with mild, moderate and severe meningococcal sepsis. Using Western blot, we detected HLPs in 11 of 20 patients with meningococcal disease but not in any of 10 patients with sepsis of other bacterial aetiologies or healthy controls **(Fig. 3A)**. There was substantial heterogeneity in HLP levels between cases that could be related to the time of collection during the hospitalisation. While there was a trend toward increased IHFβ levels in severe disease, this was not observed for hupB, for which only low plasma levels were detected regardless of group. These findings indicate that circulating HLPs are detectable in at least some patients with meningococcal disease, but this distal detection may not accurately reflect what is occurring at sites of vascular injury.

**Figure 3.**
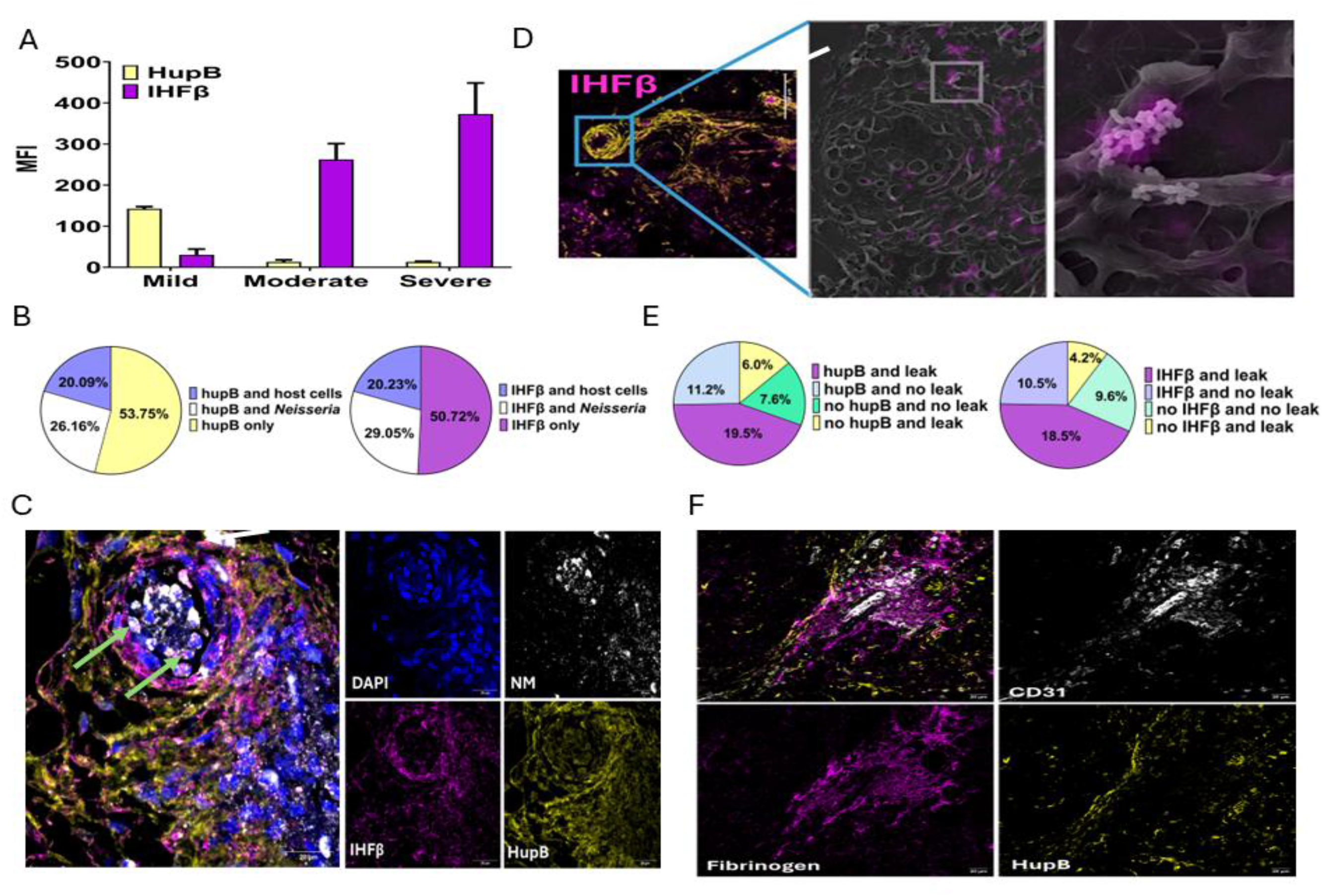
Detection of HLPs in plasma and in purpura fulminans lesions in association with pathological features. **(A)** HLPs are detectable in plasma of the patients with meningococcal disease (see Methods), **(B)** HLPs co-localise with *Neisseria meningitidis*, host cells and can be detected extracellularly, the distribution of staining is presented as pie charts, **(C)** HLPs co-localise with *Neisseria meningitidis*, host cells and can be detected extracellularly with white arrow showing cocci co-localised with IHFβ while a green arrow points extracellular IHFβ s in the blood vessel lining, **(D)** HLPs co-localisation was further confirmed using correlative light-electron microscopy (CLEM), an example of co-localisation with *N. meningitidis* and extracellularly (IHF-β marked in magenta), **(E)** HLPs co-localise with vascular leak (presence of fibrinogen outside of the blood vessel labelled by (CD31) with HupB presented as an example **(F)**. Briefly, the number of blood vessels (CD31 positive) were counted in each section, followed by counting blood vessels with leak present (fibrinogen outside the blood vessel) and its co-localisation with extracellular HLPs. Data presented as pie charts or contingency tables where appropriate. A pre-specified alpha level of 5% (P<0.05) was considered as significant.

To more directly investigate the association between HLPs and vascular injury, we examined their presence in skin punch biopsies taken from the edge of purpura fulminans lesions in patients with meningococcal disease (n=4 for hupB; n=4 for IHFβ). Biopsies were taken from the boundary of purpuric lesions to include both affected and adjacent non-purpuric skin. These were compared with contemporaneous skin punch biopsies from healthy control patients (n=2). In meningococcal patient skin, we detected substantial levels of the three meningococcal HLPs using custom anti-HLP antibodies. HLP levels were higher in diseased areas than in adjacent normal skin **(Fig. 3)**, and no HLPs were detected in control samples **(fig. S6)**.

The detection of HLPs inside bacteria was expected, as they are constitutively expressed. However, for HLPs to plausibly contribute to pathogenesis, they must also be present extracellularly. To objectively quantify and localize intracellular and extracellular HLPs, we developed an automated image analysis pipeline. As expected, HLPs co-localized with *N. meningitidis* in tissue biopsies (26% hupB and 29% IHFβ). Additionally, HLPs were associated with host cell nuclei (20% hupB and 20% IHFβ) **(Fig. 3B)** and were also detected in tissue without clear association with either meningococci or host nuclei. The predominance of extracellular HLPs may reflect that these patients had all received antibiotics, resulting in bacterial death and degradation and HLP release, findings consistent with our *in vitro* data showing HLP release with antibiotic treatment **(Fig. 3C)**.

To confirm the presence of HLPs both inside and outside bacteria, we used correlative light and electron microscopy (CLEM), which overlays fluorescence staining with electron microscopy images, allowing precise identification of bacterial and host cells.

Meningococcal structures were clearly visible within blood vessels, and HLPs were seen co-localising with both adherent bacteria and host cells **(Fig. 3D)**. We next assessed whether the presence of HLPs was spatially associated with vascular leak. This was evaluated by staining for fibrinogen, which remains intravascular in intact vessels and becomes extravascular when vessel integrity is compromised. In control cases and in non-purpuric areas of meningococcal patient skin, fibrinogen extravasation was minimal. However, in purpuric areas, fibrinogen was detected outside blood vessels, indicating vascular leak. Notably, extracellular IHFβ was significantly more abundant in areas of fibrinogen extravasation compared to areas without leak, suggesting co- localisation (χ2, p=0.04), whereas extracellular HupB levels showed no significant associations with the areas of leak (χ2, p=0.2) **(Fig. 3E)**.

### Non-anticoagulant heparins and polyclonal antibodies attenuate HLPs mediated barrier disruption

There is clear evidence for efficacy of heparins in multiple animal models, shown to be in part through their ability to bind host extracellular histones and reduce their toxic and procoagulant effects (*33*). However, the translation of this principle into outcome improvements in patients in clinical trials have been mixed, with some showing significant mortality reduction (*34, 35*) and others showing no clear efficacy. In meningococcal sepsis specifically, a small study showed no benefit but was underpowered to do so (26 patients total, 11 receiving heparin) (*36*). Severe bleeding is a recognised complication and was significantly increased in some studies (*37*). The development of several non-anticoagulant heparin variants which do not confer a bleeding risk but retain endothelial protective effects including the ability to neutralise histones provide an opportunity, but these are yet to be tested for sepsis in randomised controlled trials.

Since HLPs are also highly positively charged, we reasoned that non-anticoagulant heparins may attenuate HLPs effects and, if so, might represent a safe and plausible option for treatment. Thus, we preincubated supernatants and pellets from both bacterial strains or recombinant HLPs with either commercially available heparan sulphate (HS), N-Acetylheparin (HEP) or a panel of non-anticoagulant heparins provided by IntelliHep (Hp-A-D) to determine whether reduction of endothelial barrier permeability can be achieved.

Heparan sulphate significantly attenuated barrier disruption for bacterial supernatants as well as for the recombinant hupB and IHFβ, but not for IHFα. This heterogeneity, in the ability of different modified heparins to attenuate barrier disruption between bacterial supernatants **(Fig. 4A)** and recombinant HLPs **(Fig. 4B)**, is indicative that the degree of acetylation and sulphonylation may affect HLP neutralising capacity. We also tested the potential of a mixture of the three affinity-purified anti-HLP polyclonal antibodies to modify the effect of bacterial supernatant and recombinant HLPs on endothelial barrier: the antibody mixture significantly attenuated barrier disruption for bacterial supernatant (**Fig. 1C**, **Fig. 4A)** and completely reversed barrier disruption for the three rHLPs **(Fig. 4B)**.

**Figure 4.**
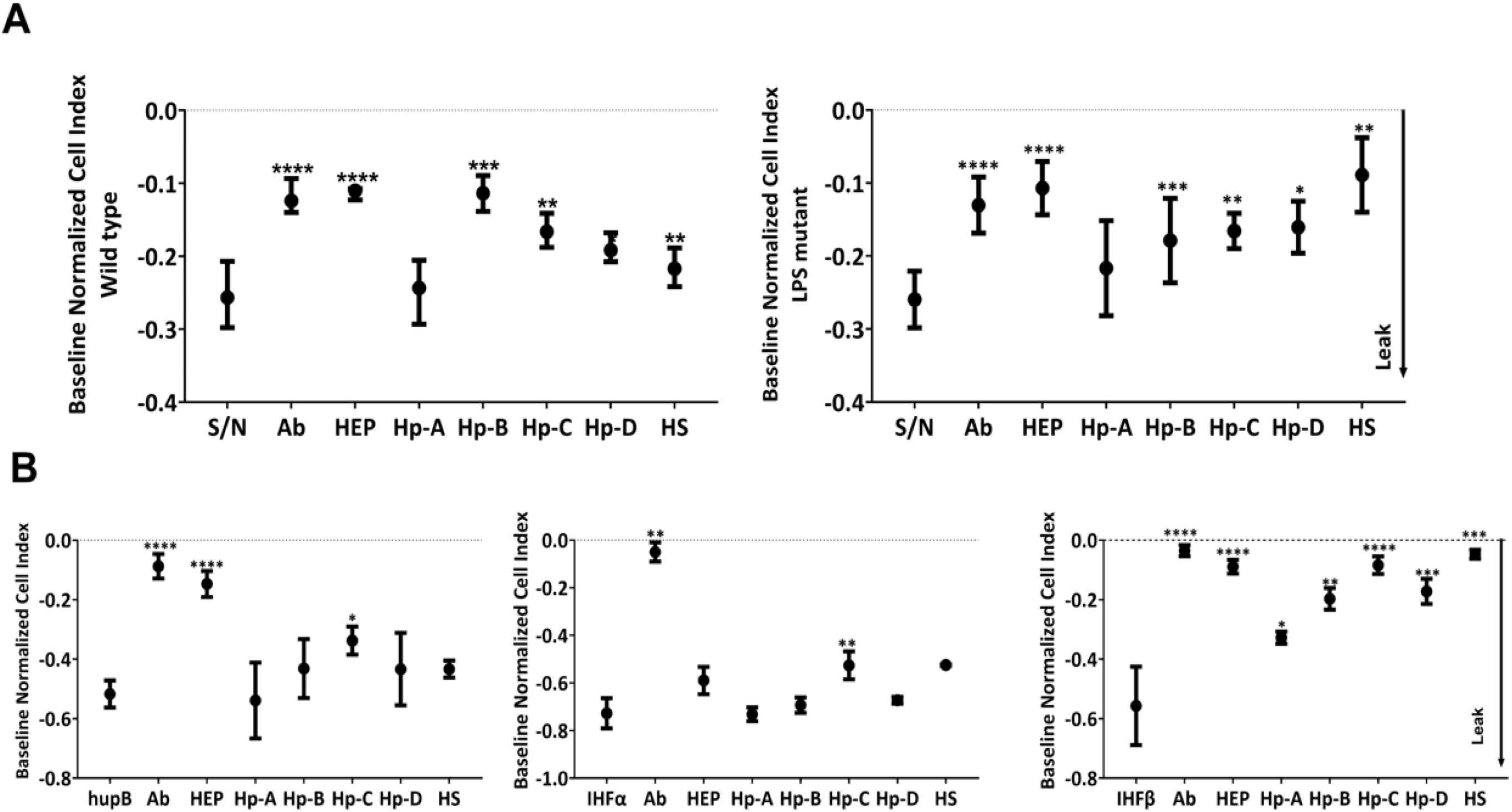
Pre-incubation with non-anticoagulant heparins or polyclonal antibodies to HLPs reduces endothelial monolayer leak. **(A)** Endothelial monolayer leak induced by meningococcal supernatants after Ceftriaxone treatment was reduced by preincubation with non-anticoagulant heparins, glycosaminoglycans, or HLP depletion **(B)** rHLPs (hupB, IHFα and IHFβ) were preincubated with non-anticoagulant heparins before addition to the monolayer of endothelial cells. S/N - supernatant; Ab - HLPs pulldown with anti-HLP antibodies, HEP (N-Acetylheparin sodium salt) and four modified with non-anticoagulant heparins (Hp-A to Hp-D produced by Intellihep Ltd. HS-heparan sulphate. *P < 0.05, **P < 0.01; ****P < 0.0001 statistically significant. Experiments were performed in triplicate. Data presented as Mean with SD.

### HLPs from other bacterial species induce variable endothelial barrier disruption

Given significant conservation in some HLPs between bacterial species, we hypothesised that HLPs might function as virulence factors in multiple pathogenic bacteria. To assess this, we generated custom recombinant HU proteins (hup) from two Gram-positive bacteria that are clinically important causes of sepsis: *Streptococcus pyogenes* serotype M3 (hupB) and *Staphylococcus aureus* strain COL (hup). Both HU proteins induced dose-dependent reductions in endothelial impedance **(fig.S7A)**. However, HU from *S. pyogenes* was significantly more potent than *S. aureus*, suggesting that despite their conservation, different HLPs vary in their capacity to disrupt the endothelium. For *S. pyogenes* HU, the barrier-disruptive effect was fully reversed following the application of N-acetylheparin (HEP; a widely commercially available non-anticoagulant heparin). In contrast, HEP had no effect on the endothelial disruption caused by *S. aureus* HU, suggesting that modifications to the non-anticoagulant heparin structure affect its ability to neutralize the disruption **(fig. S7B)**.

### Histone-like proteins (HLPs) are toxic *in vivo*

Next, we aimed to assess the effect of HLPs in vivo. We used larvae of the Greater Wax Moth, *Galleria mellonella*, which shares conserved innate immune responses with humans and has been used as an *in vivo* model to study the pathogenesis and virulence factors of several bacterial and fungal human pathogens (*38, 39*). For initial screening, we used bacterial supernatants and recombinant proteins at barrier-disruptive concentrations, as determined by the in vitro impedance assay (xCELLigence). Supernatants from ceftriaxone treated and untreated *N. meningitidis* wild type and endotoxin-lacking LPS-mutant strains, were injected into the haemolymph cavity of *Galleria mellonella* larvae. Survival was monitored for six days post-injection (**Fig. 5A**). Melanisation was monitored as an indication of coagulation activation. Reduced larval survival **(Fig. 5B-C)** and increased melanisation **(Fig. 5D)** was observed following injection of antibiotic-treated supernatants. To assess whether recombinant HLPs alone were toxic *in vivo*, a range of HLP concentrations (3.13–50 µg/ml) were tested. A dose-dependent reduction in larval survival was observed across all HLPs including three meningococcal HLPs and the two HU proteins from *S. aureus* (hup) and *S. pyogenes* (hupB, **fig. S8**). These results further support our *in vitro* findings, suggesting that HLPs function as potential virulence factors and promote coagulation.

**Figure 5.**
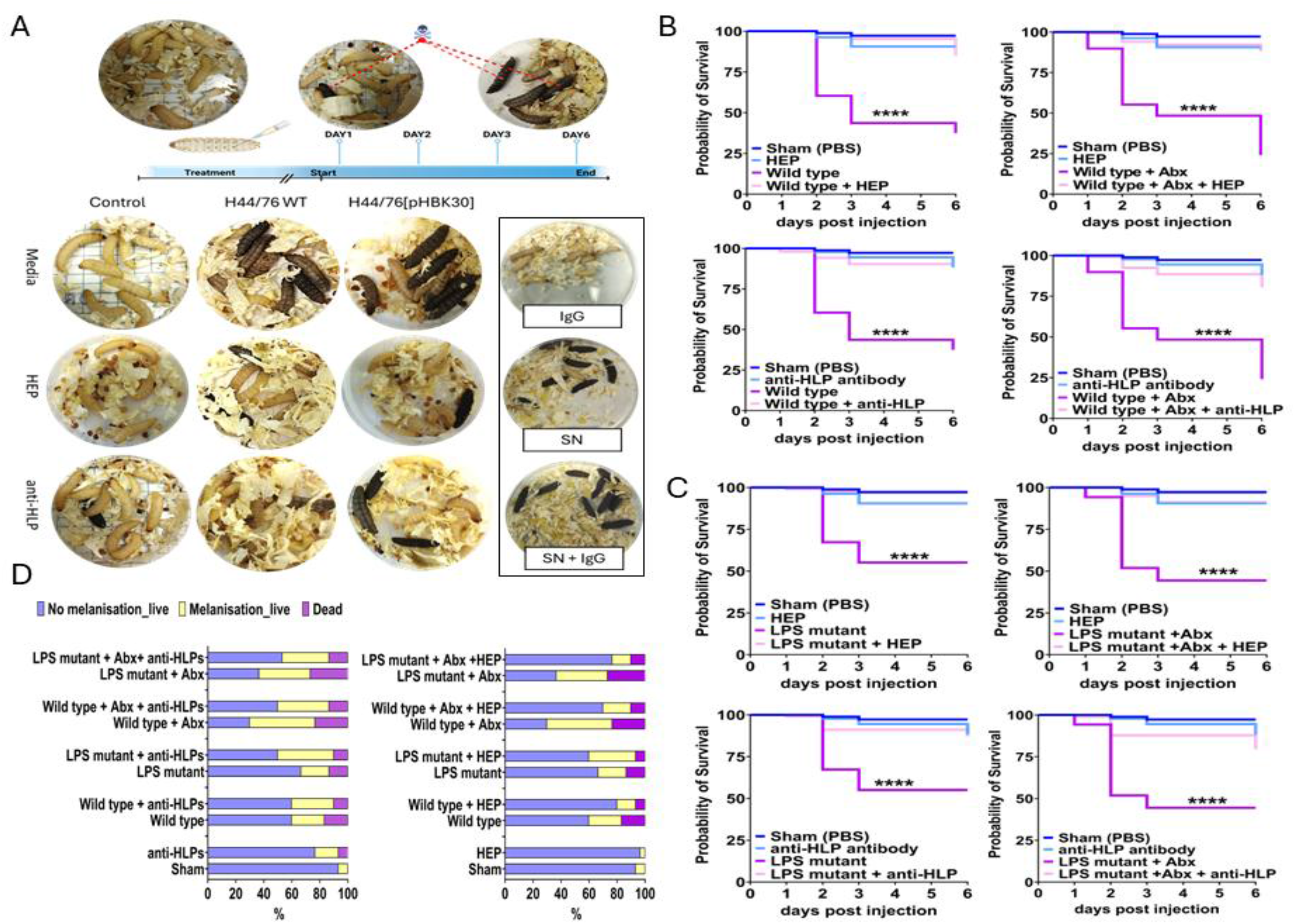
Supernatants from *N*. *meningitidis* cultures and recombinant HLPs reduce *Galleria mellonella* larvae survival. **(A)** An example of outcome (larvae survival) observed after injection of *N.meningitidis* supernatants (SN) from the wild-type (H44/76) and LPS mutant (H44/76[pHBK30]) strains alone or after pre-incubation with neutralising agents (non-anticoagulant N-Acetylheparin (HEP) or anti-HLP antibodies cocktail). The supernatants from wild type **(B)** and LPS-mutant **(C)** meningococci strains after Ceftriaxone treatment were toxic and reduced larvae survival, this effect was reversed or reduced by pre-incubation with neutralising agents, negative IgG control **(D)** Supernatants neutralisation affects larvae melanisation. Asterisks indicate significant differences relative to the HLP treatment group and groups pre-treated with non-anticoagulant HEP or the anti-HLP antibodies cocktail, as assessed by the log-rank (Mantel–Cox) test (ns, non-significant, *P < 0.05, **P < 0.01; ****P < 0.0001). Data were presented as mean with SD.

### Modified heparins and polyclonal antibodies reduce HLP-induced toxicity *in vivo*

To assess whether non-anticoagulant heparins or polyclonal anti-HLP antibodies neutralise HLP-induced toxicity in the *G. mellonella* larvae model, meningococcal supernatants from wild type and LPS-mutant strain were preincubated with N-acetylheparin (HEP) or the cocktail of three anti-HLP antibodies prior to injecting them into the larvae. Larvae survival and melanisation was substantially improved with both treatments of larvae survival and reduced melanisation, demonstrating that HLPs and/or HLPs complexes can be successfully neutralised *in vivo* (**Fig. 5)**. To assess whether heparin structure influences the degree of neutralisation of meningococcal supernatants, we preincubated a panel of non-anticoagulant heparins with meningococcal supernatants prior to injection into the larvae. Overall, supernatant toxicity was neutralised, as reflected by improved larval survival. However, there were differences in the response between wild type and LPS mutant strains as well as melanisation degree **(Fig. 6B, C).** Additionally, some of the tested heparins affected the rate of pupae formation—for example, Hp-A accelerated pupation, while Hp-C delayed it. These effects warrant further investigation but fall outside the scope of this study.

**Figure. 6.**
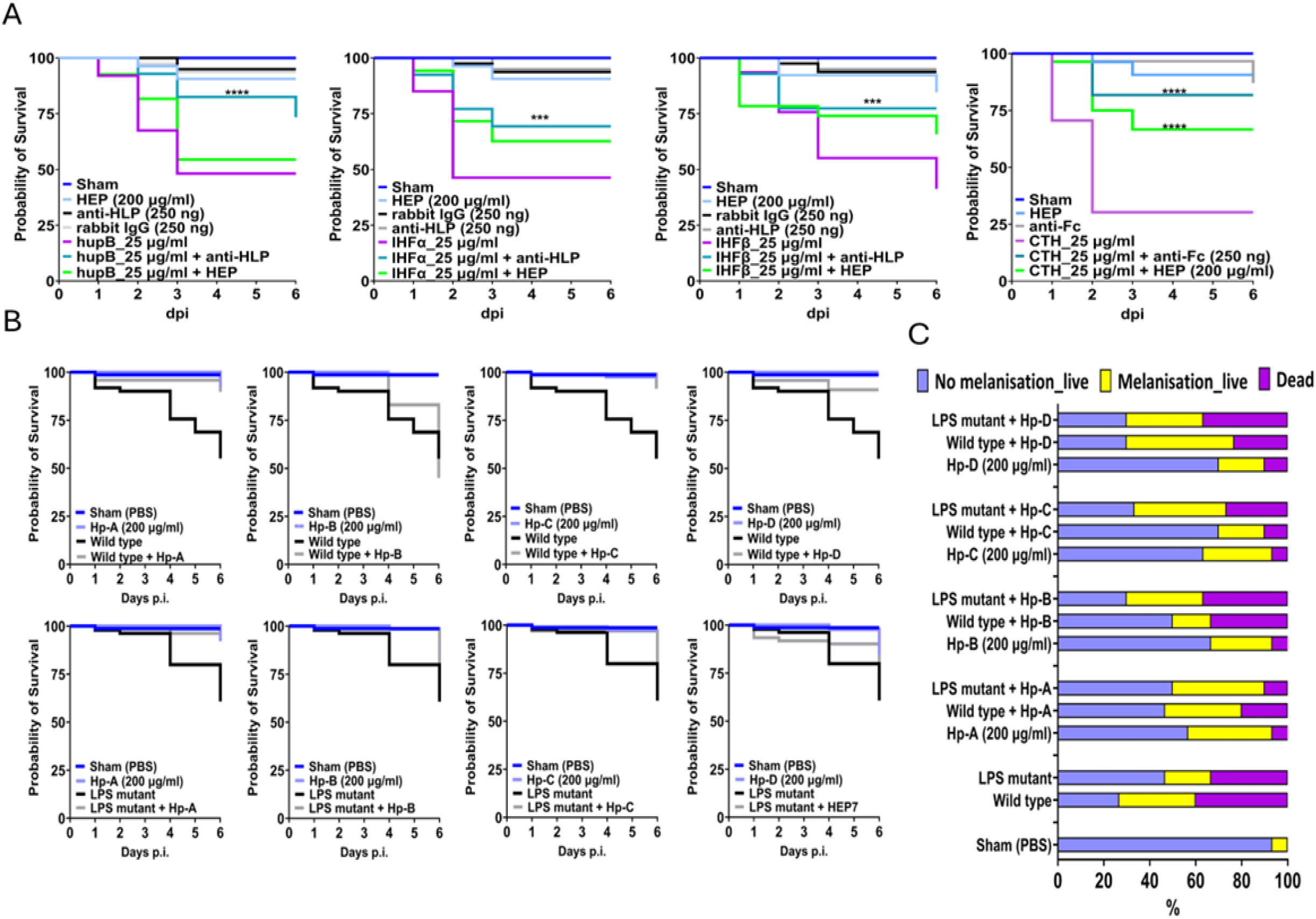
The effect of neutralisation of HLPs on *Galleria mellonella* larvae survival and melanisation. **(A)** The effect of rHLPs on larvae is dose-dependent and can be reversed by non-anticoagulant heparin (HEP) or cocktail of the anti-HLP antibodies. As positive control, calf thymus histones (CTH) were used with anti-Fc histone, **(B)** Preincubation of non-anticoagulant heparins (Hp-A-Hp-D) with bacterial supernatants from wild type and LPS mutant strains improved larvae survival, **(C)** Larvae melanisation was modulated by non-anticoagulant heparins from Intellihep. Hp-A to Hp-D. Larvae survival was assessed by the log-rank (Mantel–Cox) test (ns, non-significant, *P < 0.05, **P < 0.01; ****P < 0.0001). Data were presented as mean with SD.

The toxic effects of recombinant HLPs (25 µg/ml) could also be reduced by preincubation with either non-anticoagulant heparin or anti-HLP antibodies **(Fig. 6A).** This effect was specific to anti-HLP antibodies and not a non-specific effect as a rabbit IgG treatment at the same concentration did not reduced toxicity **(fig. S9)**. HU protein from *Staphylococcus aureus* and *Streptococcus pneumoniae* also induced a dose dependent reduction in larvae non-anticoagulant heparins reduced toxicity and improved larvae survival for exposure to *Streptococcus pneumoniae* hupB but not for *Staphylococcus aureus* hup suggesting that neutralisation efficiency may depend on the heparin structure **(fig. S10)**.

## Discussion

In this study, we demonstrated that histone-like proteins (HLPs) are released from *N. meningitidis* on antibiotic treatment, and are a potential virulence factor with endothelial disruptive, pro-coagulant properties that contributed to fatal outcome in a larvae *in vivo* model. These adverse effects were attenuated both *in vitro* and *in vivo* by non-anticoagulant heparin or a cocktail of anti-HLP antibodies. Data from patients with meningococcal sepsis further supported a role of HLPs in disease pathology, with HLPs detectable in plasma and in skin biopsies from patients with severe disease. In skin biopsies, HLPs were found outside of bacteria, co-localised with host cells, and enriched in areas of vascular leak and proximity to NETs. While these findings are associative and do not establish causality, taken together with our *in vitro* and *in vivo* data, they provide a compelling link to vascular injury in patients. Significant barrier disruption *in vitro* and toxicity *in vivo* from HLPs from virulent Gram-positive bacteria *Staphylococcus aureus* and *Streptococcus pyogenes* highlight that HLPs may be involved in the pathogenesis of other bacteria, with the potential to have broad therapeutic benefits for bacterial infections. HLPs also cause endothelial barrier disruption and are toxic *in vivo* and HLPs are relatively conserved between species the question arises: why do HLPs cause severe illness and toxicity when released from some bacteria but not for others, such as the bacteria that constitute the gut flora? While a definitive answer to this requires further investigation, several explanations can be proposed. Firstly, there are variations in toxicity among HLPs originating from different bacterial strains tested in our system. This may be due to post-translational modifications that could affect the overall charge of the HLPs and/or add additional steric bulk. Secondly HLPs only induced barrier disruption once a threshold dose was reached, suggesting that the total and tissue level quantities of HLPs may be critical determinants of pathogenicity. These may be influenced by the degree of HLP production as HLP expression is high in rapidly growing bacteria, typical in bacteraemic sepsis, and HLP production is low in bacteria that are dormant or in stable colonies such as commensal bacteria in the gut (*27, 40–42*). Notably in meningococcal sepsis bacterial loads are typically greater than 1×10^7^ CFU/ml and reach up to 1×10^9^ CFU/ml ml (*43*). Additionally, the sites of bacterial adhesion may affect local concentrations of HLP release in different tissue microenvironments, such as the concentrated adhesion of Meningococci to the endothelial surface. Further research is needed to assess the role and relative importance of these different variables.

We explored the therapeutic potential of both non-anticoagulant heparins and anti-HLP antibodies. Notably, a library of modified non-anticoagulant heparins with varying levels of acetylation and sulphonylation exhibited heterogenous responses to both antibiotic-killed bacteria and their culture supernatant, as well as to different recombinant HLPs. Thus, to achieve optimal therapeutic response to heparin it may be necessary to tailor its structure. Another therapeutic approach involves generating antibodies or a limited antibody cocktail targeting HLPs (*44*). For this to be effective, it would be necessary to identify an HLP epitope (or epitopes) essential for their toxicity, which is conserved across different HLPs and does not cross-react with host proteins. This could be designed *in silico* based on experimental data or possibly by cloning monoclonal antibodies from B cells of convalescent patients who have recovered from meningococcal disease.

This study has several limitations. While we have demonstrated the toxic and endothelial disruptive properties of HLPs *in vitro* and *in vivo* and shown evidence from patients that they concentrate at sites of vascular damage, we have not proven their causal role in disease. While we have used the Greater Wax Moth larvae as a model of HLP toxicity that could further be used to investigate innate immune mechanisms, it is not a full model of bacterial infection. *N. meningitidis* sepsis lacks an animal model that closely mimic the diseases, but further work using other bacterial infections in a mammalian model of infections that reach high bacterial loads would be useful as proof of concept and for additional pre-clinical studies.

Here we have shown that HLPs are a potential virulence factor, released from bacteria during antibiotic treatment, promoting endothelial activation, procoagulant phenotype and NETs formation, and that their effect can be attenuated by antibody or by non-anticoagulant heparins. These findings indicate possible routes to a therapy that could be administered concurrently with the first dose of antibiotics. Moreover, the fact that HLPs from other pathogenic bacteria are also toxic indicates value in exploring their effect in other bacterial infections, with the potential for targeting them as a much-needed broad spectrum adjunctive treatment for bacterial sepsis.

## Material and methods

### Neisseria meningitidis culture

*N. meningitidis* serogroup B H44/76 (wild type) and an LPS-mutant strain H44/76[pHBK30], deficient in lipid A biosynthesis, were used. The mutant strain has been previously shown to lack detectable LPS or endotoxin (*32*). LPS deficiency was verified by sequencing and endotoxin test (LAL assay; **fig. S3**). Bacteria were grown to mid-log phase (OD_600nm_ ∼0.5) and treated with Ceftriaxone at 0.006 μg/ml (0.1x MIC), 0.066 μg/ml (MIC), or 0.6 μg/ml (10x MIC) for 3 hours (*45*). Supernatants and pellets were collected, inactivated and diluted 5x for subsequent experiments.

### Endothelial Cell culture

Primary human dermal microvascular endothelial cells (HDMEC; PromoCell®GmbH, Germany ) were cultured routinely, on attachment factor coated cell culture dishes in Endothelial Cell Growth MV medium with supplement mix (EGM-MV; PromoCell®). Cells were passaged using Accutase® (ThermoFisher) following manufacturer instructions and routinely checked for mycoplasma (MycoStrip^TM^, InvivoGen)

### Plasma samples and skin biopsies

Plasma samples from the EUCLIDS study were randomly selected based on disease severity, including severe non-meningococcal and control groups (N=10 each) (*46*).

Punch biopsies of skin (3mm) were obtained from patients with meningococcal sepsis confirmed by clinical and microbiological criteria described in detail previously. Samples were taken from the edge of purpura fulminans lesions and adjacent non-purpuric skin. Control biopsies were collected during routine surgery. Samples were formalin-fixed, and paraffin embedded (FFPE). The study was approved by the St. Mary’s Hospital research ethics committee and the UK national Integrated Research Application System (IRAS 263175). Informed parental consent was obtained for all paediatric participants.

### Recombinant Histone like proteins (HLPs)

Neisseria meningitidis serogroup B was chosen to generate full length recombinant HLPs namely DNA-binding HU beta (hupB); Integration host factor subunit alpha (IHFα) and Integration host factor subunit alpha (IHFβ) which were custom manufactured for this project by Cusabio (Houston, USA). The proteins were N-terminally His-tagged and affinity purified using Immobilized metal ion affinity chromatography (IMAC) with >92% purity with observed for hupB and IHFβ with band sizes 16kDa and 18kDa, respectively and >85% for IHFα band sizes 16kDa. Recombinant proteins were processed aseptically before the lyophilisation and tested for endotoxin levels (< 1 EU per 1μg of the protein by the LAL method) and confirmed in house as < 0.1 EU per 1 μg.

### Polyclonal Anti-HLP antibodies

Antibodies were custom manufactured for this project by Cusabio (Houston, USA). Recombinant full length HupB, IHFα or IHFβ with N- terminal GST-tag expressed in *E.coli* were used as an antigen to raise specific rabbit polyclonal antibodies recognising full length HupB, IHFα or IHFβ. Polyclonal antibodies were affinity purified (>95% purity) and underwent QA&QC including antibody titration, testing for specific applications including Western blot and ELISA. Additionally, in house antibody validation was performed.

### *Galleria mellonella* (Greater Wax Moth) larvae model

*Galleria mellonella* larvae were obtained from Livefood UK, kept in darkness at room temperature and were used up to 1 week following arrival. Larvae were injected with bacterial supernatants, recombinant HLPs (hupB, IHFα, IHFβ) alone, or preincubated with non- anticoagulant heparins or anti-HLP antibodies. 10 µL of sample was injected in the haemocoel via the last right prolimb, afterwards larvae were maintained in darkness at 37⁰C and their survival monitored across 6 days post-injection (N=10 larvae without signs of melanisation per group, 3 biological replicates). Inactive, black larvae were recorded as dead. Both positive and negative controls were run with each experiment as appropriate (sham control, antibiotic alone, non-anticoagulant heparins, anti-HLP antibodies and rabbit IgG).

### Endothelial assays

Endothelial barrier function was assessed by transendothelial electrical resistance (TEER xCELLigence RTCA) and expressed as cell index (CI). Changes in the endothelial barrier function were normalised to baseline (control) and reported as baseline normalised cell index (BNCI). HDMECs were grown overnight until confluency (CI ≥10), then incubated for 2h in OptiMEM medium containing 1% KnockOut™ Serum Replacement (ThermoFisher) prior to treatment (**table SM1)**.

Cell viability was assessed using Apotracker™Green (BioLegend), Live-or-Dye™594/614 (Biotium) and Hoechst 33342 (ThermoFisher). Confluent HDMECs cells treated as above, with PMA (5 nM), TNFα (100 ng/ml), or Staurosporine (1 μM) as positive controls. Each assay was performed in triplicate (5 fields being used for analysis per technical duplicate). QuPath software was used to evaluate endothelial cells viability (*47*).

Immunofluorescence was used to detect junctional proteins (VE-cadherin, EPCR, ICAM-1) and bacterial HLPs (hupB, IHFα and IHFβ). Images were acquired on EVOS FL2 and analysed using QuPath software.

In-cell western assays were used to quantify junctional proteins (VE-cadherin, EPCR, ICAM-1) normalised to the total cell number per well using Odyssey®CLx Imaging System. Data were analysed and presented as a ratio of target signal normalised to cell number.

Detailed description in the Supplementary Methods (SM).

### Western blot

Reduced plasma samples (3ul) were separated on 12% Bis-Tris gels in MES-SDS and transferred onto Immobilon-FL. Membranes were blocked, incubated overnight at 4°C with primary antibodies against hupB, IHFα, or IHFβ), then with fluorescent secondary antibodies for 1 h at room temperature. Blots were visualised using Odyssey CLx imaging system and analysed using Image Studio software (LI-CORE, UK). Detailed conditions are provided in the Supplementary Methods (SM).

### Histology and Correlative light-electron microscopy (CLEM)

FFPE skin sections (3μm) from paediatric meningococcal patients and controls were stained using a multiplex Tyramide Signal Amplification (Opal-TSA) system (Akoya Biosciences). Sections were dewaxed, rehydrated, subjected to heat-induced antigen retrieval, and blocked before overnight incubation with primary antibodies (4°C). Detection was performed using HRP-conjugated secondary antibodies and Opal fluorophores. Detailed conditions for every stage are provided in the Supplementary Methods and **table SM2**). Confocal images were captured using a Nikon AF2 microscope and analysed independently by two analysts using QuPath and CellProfiler (*48*) by 2 independent analysts.

CLEM was performed to examine co-localization of HLPs and *Neisseria* within meningococcal tissue. Confocal images were correlated with scanning electron microscopy (JEOL IT-100, Japan). Image overlays were generated in Adobe Photoshop, and automated quantification pipelines were developed in CellProfiler.

Analysis scripts are available at (https://github.com/llemgruber/Cellular_Analysis_Facility.git)

### Statistical analysis

Statistical analysis was preformed using GraphPad Prizm v.10. ANOVA or Chi-square tests were applied as appropriate. Kaplan–Meier survival curves were compared using log-rank (Mantel–Cox) tests. P<0.05 was considered significant. ns, non-significant; ****P< 0.0001; ***P< 0.001, **P<0.01; *P<0.05).

## List of the Supplementary materials

**figure S1.** Histone like proteins (HLPs) are detectable in supernatants and pellets from *Neisseria meningitidis* cultures after treatment with Ceftriaxone using Western blot.

**figure S2.** The pellets from *Neisseria meningitidis* cultures after treatment with Ceftriaxone affect human primary dermal endothelial cells (HDEC) permeability, however this effect is not dose related.

**figure S3. (A)** Comparison of LPS locus between used bacterial strains. (B) Detection of endotoxin levels in the *Neisseria meningitidis* culture supernatants.

**figure S4.** *Neisseria meningitidis* culture components activate endothelial cells.

**figure S5.** DNase I treatment doesn’t have effect on recombinant histone like proteins (HLPs) mediated HDECs permeability.

**figure S6. (A)** Examples of staining for *Neisseria meningitidis* and HLPs in control skin punch biopsies. **(B)** Examples of staining for Fibrinogen and HLPs in control skin punch biopsies.

**figure S7**. Endothelial cells permeability is affected by hup from Staphylococcus aureus (SA) and hupB from Streptococcus pyogenes (SP).

**figure S8.** The effect of recombinant *Neisseria meningitis* HLPs (HupB, IHFα and IHF β), Hu proteins from Staphylococcus aureus (SA, hup) and Streptococcus pyogenes (SP, hupB) on *Galleria mellonella* larvae survival is dose dependent.

**figure S9.** Presence of rabbit IgG did not reduce supernatant toxicity or improved *Galleria mellonella* larvae survival.

**figure S10.** The effect of non-anticoagulant N-Acetylheparin (HEP, 200 μg/ml) or cocktail of the anti-HLP antibodies (250 ng) on the on *Galleria mellonella* larvae survival after the treatment with the recombinant Hu proteins from Staphylococcus aureus (SA) and Streptococcus pyogenes (SP).

**Supplementary methods (SM)** contain detailed protocols for techniques and conditions used in the study.

**table SM1**. List of compounds and working concentrations used in the study.

**table SM2.** Antibody information and working concentration, antigen retrieval methods and other information regarding IHC.

**Movies S1 to S3.** Projection of HLPs **(S1)** HupB, **(S2)** IHFα and **(S3)** IHFβ_ localisation (yellow) in HDECs (video) VE-cadherin staining in red colour, DAPI (blue). 40x magnification

## Acknowledgements

**Funding:** This work was funded through a grant from Chief Scientist Office (TCS/19/27) [CM] and Medical Research Council (IAA STTAR-BacPro) [CM/DMcG].

RSH and SNF are NIHR Senior Investigators. RSH was supported by the National Institute for Health and Care Research University College London Hospitals Biomedical Research Centre. The authors gratefully acknowledge the Cellular Analysis Facility for their support & assistance in this work.

## Author contribution

Conceptualization (CAM, DMcG, ML, RSH); Funding acquisition (CAM, DMcG); Writing – original draft (DMcG, CAM); Investigation (DMcG, CJ); Methodology: DMcG (in vitro and in vivo models, HTC imaging) CJ (mIF confocal microscopy), LLS (electron microscopy, slide processing for EM); Resources: Plasma and tissue sections and clinical advice (JH, VJW, RN, RSH, SNF, SNF); synthesis and provision of non-anticoagulant heparins (EAY&JT), generating lysates from wild type and mutant meningococci (FB), microbiology expertise and resources (TE, RMcH, GD, AJR, AS, RU, C-HT, SA,FB), Visualization: imaging and image processing (CJ, DMcG, LLS); Writing – review & editing (EAY, JT, TE, RMcH, FB, GD, AJR, AS, RU, C-HT, SA, JH, RN, RSH, SNF, ML), Supervision (CAM)

## Data availability

All data associated with this study are available in the main text or the supplementary material

## References

1. 1. Sepsis (available at https://www.who.int/news-room/fact-sheets/detail/sepsis).

2. S. N. Faust, M. Levin, O. B. Harrison, R. D. Goldin, M. S. Lockhart, S. Kondaveeti, Z. Laszik, C. T. Esmon, R. S. Heyderman, Dysfunction of endothelial protein C activation in severe meningococcal sepsis. N Engl J Med 345, 408–416 (2001).

3. E. Nectoux, A. Mezel, S. Raux, D. Fron, M. Maillet, B. Herbaux, Meningococcal purpura fulminans in children: I. Initial orthopedic management. J Child Orthop 4, 401–407 (2010).

4. P. Brandtzaeg, P. Kierulf, P. Gaustad, A. Skulberg, J. N. Bruun, S. Halvorsen, E. Sørensen, Plasma endotoxin as a predictor of multiple organ failure and death in systemic meningococcal disease. J Infect Dis 159, 195–204 (1989).

5. J. Dick, S. Hebling, J. Becam, M.-K. Taha, A. Schubert-Unkmeir, Comparison of the inflammatory response of brain microvascular and peripheral endothelial cells following infection with Neisseria meningitidis. Pathog Dis 75 (2017), doi:10.1093/femspd/ftx038.

6. V. Aberg, F. Almqvist, Pilicides-small molecules targeting bacterial virulence. Org Biomol Chem 5, 1827–1834 (2007).

7. K. Denis, M. Le Bris, L. Le Guennec, J.-P. Barnier, C. Faure, A. Gouge, H. Bouzinba-Ségard, A. Jamet, D. Euphrasie, B. Durel, N. Barois, P. Pelissier, P. C. Morand, M. Coureuil, F. Lafont, O. Join-Lambert, X. Nassif, S. Bourdoulous, Targeting Type IV pili as an antivirulence strategy against invasive meningococcal disease. Nat Microbiol 4, 972–984 (2019).

8. K. J. Dye, N. J. Vogelaar, M. O’Hara, P. Sobrado, W. Santos, P. R. Carlier, Z. Yang, Discovery of Two Inhibitors of the Type IV Pilus Assembly ATPase PilB as Potential Antivirulence Compounds. Microbiol Spectr 10, e0387722 (2022).

9. D. C. Morrison, J. L. Ryan, Endotoxins and disease mechanisms. Annu Rev Med 38, 417–432 (1987).

10. M. Fischer, J. Hilinski, D. S. Stephens, Adjuvant therapy for meningococcal sepsis. Pediatr Infect Dis J 24, 177–178 (2005).

11. B. Derkx, J. Wittes, R. McCloskey, Randomized, placebo-controlled trial of HA-1A, a human monoclonal antibody to endotoxin, in children with meningococcal septic shock. European Pediatric Meningococcal Septic Shock Trial Study Group. Clin Infect Dis 28, 770–777 (1999).

12. B. P. Giroir, P. J. Scannon, M. Levin, Bactericidal/permeability-increasing protein--lessons learned from the phase III, randomized, clinical trial of rBPI21 for adjunctive treatment of children with severe meningococcemia. Crit Care Med 29, S130–135 (2001).

13. G. R. Bernard, J. L. Vincent, P. F. Laterre, S. P. LaRosa, J. F. Dhainaut, A. Lopez-Rodriguez, J. S. Steingrub, G. E. Garber, J. D. Helterbrand, E. W. Ely, C. J. Fisher, Recombinant human protein C Worldwide Evaluation in Severe Sepsis (PROWESS) study group, Efficacy and safety of recombinant human activated protein C for severe sepsis. N Engl J Med 344, 699–709 (2001).

14. V. M. Ranieri, B. T. Thompson, P. S. Barie, J.-F. Dhainaut, I. S. Douglas, S. Finfer, B. Gårdlund, J. C. Marshall, A. Rhodes, A. Artigas, D. Payen, J. Tenhunen, H. R. Al-Khalidi, V. Thompson, J. Janes, W. L. Macias, B. Vangerow, M. D. Williams, PROWESS-SHOCK Study Group, Drotrecogin alfa (activated) in adults with septic shock. N Engl J Med 366, 2055–2064 (2012).

15. P. M. Lepper, T. K. Held, E. M. Schneider, E. Bölke, H. Gerlach, M. Trautmann, Clinical implications of antibiotic-induced endotoxin release in septic shock. Intensive Care Med 28, 824–833 (2002).

16. J. L. Halbach, A. W. Wang, D. Hawisher, D. M. Cauvi, R. E. Lizardo, J. Rosas, T. Reyes, O. Escobedo, S. W. Bickler, R. Coimbra, A. De Maio, Why Antibiotic Treatment Is Not Enough for Sepsis Resolution: an Evaluation in an Experimental Animal Model. Infect Immun 85, e00664–17 (2017).

17. A. Sabnis, A. M. Edwards, Lipopolysaccharide as an antibiotic target. Biochim Biophys Acta Mol Cell Res 1870, 119507 (2023).

18. A. L. Erwin, Antibacterial Drug Discovery Targeting the Lipopolysaccharide Biosynthetic Enzyme LpxC. Cold Spring Harb Perspect Med 6, a025304 (2016).

19. T. Yang, J. Peng, Z. Zhang, Y. Chen, Z. Liu, L. Jiang, L. Jin, M. Han, B. Su, Y. Li, Emerging therapeutic strategies targeting extracellular histones for critical and inflammatory diseases: an updated narrative review. Front Immunol 15, 1438984 (2024).

20. J. Xu, X. Zhang, M. Monestier, N. L. Esmon, C. T. Esmon, Extracellular histones are mediators of death through TLR2 and TLR4 in mouse fatal liver injury. J Immunol 187, 2626–2631 (2011).

21. D. Pérez-Cremades, C. Bueno-Betí, J. L. García-Giménez, J. S. Ibañez-Cabellos, F. V. Pallardó, C. Hermenegildo, S. Novella, Extracellular histones trigger oxidative stress-dependent induction of the NF-kB/CAM pathway via TLR4 in endothelial cells. J Physiol Biochem 79, 251–260 (2023).

22. X. Lv, T. Wen, J. Song, D. Xie, L. Wu, X. Jiang, P. Jiang, Z. Wen, Extracellular histones are clinically relevant mediators in the pathogenesis of acute respiratory distress syndrome. Respiratory Research 18, 165 (2017).

23. K. C. A. A. Wildhagen, M. A. Wiewel, M. J. Schultz, J. Horn, R. Schrijver, C. P. M. Reutelingsperger, T. van der Poll, G. A. F. Nicolaes, Extracellular histone H3 levels are inversely correlated with antithrombin levels and platelet counts and are associated with mortality in sepsis patients. Thromb Res 136, 542–547 (2015).

24. H. Zhang, Y. Wang, M. Qu, W. Li, D. Wu, J. P. Cata, C. Miao, Neutrophil, neutrophil extracellular traps and endothelial cell dysfunction in sepsis. Clin Transl Med 13, e1170 (2023).

25. C. A. Moxon, Y. Alhamdi, J. Storm, J. M. H. Toh, D. McGuinness, J. Y. Ko, G. Murphy, S. Lane, T. E. Taylor, K. B. Seydel, S. Kampondeni, M. Potchen, J. S. O’Donnell, N. O’Regan, G. Wang, G. García-Cardeña, M. Molyneux, A. G. Craig, S. T. Abrams, C.-H. Toh, Parasite histones are toxic to brain endothelium and link blood barrier breakdown and thrombosis in cerebral malaria. Blood Adv 4, 2851–2864 (2020).

26. C. A. Moxon, N. V. Chisala, S. C. Wassmer, T. E. Taylor, K. B. Seydel, M. E. Molyneux, B. Faragher, N. Kennedy, C.-H. Toh, A. G. Craig, R. S. Heyderman, Persistent endothelial activation and inflammation after Plasmodium falciparum Infection in Malawian children. J Infect Dis 209, 610–615 (2014).

27. B. Thakur, K. Arora, A. Gupta, P. Guptasarma, The DNA-binding protein HU is a molecular glue that attaches bacteria to extracellular DNA in biofilms. J Biol Chem 296, 100532 (2021).

28. A. Severin, E. Nickbarg, J. Wooters, S. A. Quazi, Y. V. Matsuka, E. Murphy, I. K. Moutsatsos, R. J. Zagursky, S. B. Olmsted, Proteomic analysis and identification of Streptococcus pyogenes surface-associated proteins. J Bacteriol 189, 1514–1522 (2007).

29. I. Haruta, K. Kikuchi, E. Hashimoto, H. Kato, K. Hirota, M. Kobayashi, Y. Miyake, T. Uchiyama, J. Yagi, K. Shiratori, A possible role of histone-like DNA-binding protein of Streptococcus intermedius in the pathogenesis of bile duct damage in primary biliary cirrhosis. Clin Immunol 127, 245–251 (2008).

30. D. Liu, H. Yumoto, K. Hirota, K. Murakami, K. Takahashi, K. Hirao, T. Matsuo, K. Ohkura, H. Nagamune, Y. Miyake, Histone-like DNA binding protein of Streptococcus intermedius induces the expression of pro-inflammatory cytokines in human monocytes via activation of ERK1/2 and JNK pathways. Cell Microbiol 10, 262–276 (2008).

31. L. Zhang, T. A. Ignatowski, R. N. Spengler, B. Noble, M. W. Stinson, Streptococcal histone induces murine macrophages To produce interleukin-1 and tumor necrosis factor alpha. Infect Immun 67, 6473–6477 (1999).

32. L. Steeghs, R. den Hartog, A. den Boer, B. Zomer, P. Roholl, P. van der Ley, Meningitis bacterium is viable without endotoxin. Nature 392, 449–450 (1998).

33. K. C. A. A. Wildhagen, P. García de Frutos, C. P. Reutelingsperger, R. Schrijver, C. Aresté, A. Ortega-Gómez, N. M. Deckers, H. C. Hemker, O. Soehnlein, G. A. F. Nicolaes, Nonanticoagulant heparin prevents histone-mediated cytotoxicity in vitro and improves survival in sepsis. Blood 123, 1098–1101 (2014).

34. Z.-Y. Zou, J.-J. Huang, Y.-Y. Luan, Z.-J. Yang, Z.-P. Zhou, J.-J. Zhang, Y.-M. Yao, M. Wu, Early prophylactic anticoagulation with heparin alleviates mortality in critically ill patients with sepsis: a retrospective analysis from the MIMIC-IV database. Burns Trauma 10, tkac029 (2022).

35. R. Zarychanski, A. M. Abou-Setta, S. Kanji, A. F. Turgeon, A. Kumar, D. S. Houston, E. Rimmer, B. L. Houston, L. McIntyre, A. E. Fox-Robichaud, P. Hébert, D. J. Cook, D. A. Fergusson, Canadian Critical Care Trials Group, The efficacy and safety of heparin in patients with sepsis: a systematic review and metaanalysis. Crit Care Med 43, 511–518 (2015).

36. B. Haneberg, T. J. Gutteberg, P. J. Moe, B. Osterud, B. Bjorvatn, E. H. Lehmann, Heparin for infants and children with meningococcal septicemia. Results of a randomized therapeutic trial. NIPH Ann 6, 43–47 (1983).

37. M. N. Levine, G. Raskob, S. Landefeld, C. Kearon, Hemorrhagic complications of anticoagulant treatment. Chest 119, 108S–121S (2001).

38. G. Ménard, A. Rouillon, V. Cattoir, P.-Y. Donnio, Galleria mellonella as a Suitable Model of Bacterial Infection: Past, Present and Future. Front Cell Infect Microbiol 11, 782733 (2021).

39. A. Six, K. Mosbahi, M. Barge, C. Kleanthous, T. Evans, D. Walker, Pyocin efficacy in a murine model of Pseudomonas aeruginosa sepsis. J Antimicrob Chemother 76, 2317–2324 (2021).

40. C. J. Rocco, M. E. Davey, L. O. Bakaletz, S. D. Goodman, Natural antigenic differences in the functionally equivalent extracellular DNABII proteins of bacterial biofilms provide a means for targeted biofilm therapeutics. Mol Oral Microbiol 32, 118–130 (2017).

41. P. Stojkova, P. Spidlova, Bacterial nucleoid-associated protein HU as an extracellular player in host-pathogen interaction. Front Cell Infect Microbiol 12, 999737 (2022).

42. T.-H. Huang, Y.-L. Tzeng, R. M. Dickson, FAST: Rapid determinations of antibiotic susceptibility phenotypes using label-free cytometry. Cytometry A 93, 639–648 (2018).

43. J. Pardo, K. P. Klinker, S. J. Borgert, B. M. Butler, K. H. Rand, N. M. Iovine, Detection of Neisseria meningitidis from negative blood cultures and cerebrospinal fluid with the FilmArray blood culture identification panel. J Clin Microbiol 52, 2262–2264 (2014).

44. L. A. Novotny, J. A. Jurcisek, S. D. Goodman, L. O. Bakaletz, Monoclonal antibodies against DNA-binding tips of DNABII proteins disrupt biofilms in vitro and induce bacterial clearance in vivo. EBioMedicine 10, 33–44 (2016).

45. C. C. Potts, L. D. Rodriguez-Rivera, A. C. Retchless, F. Hu, H. Marjuki, A. E. Blain, L. A. McNamara, X. Wang, Antimicrobial Susceptibility Survey of Invasive Neisseria meningitidis, United States 2012-2016. J Infect Dis 225, 1871–1875 (2022).

46. F. Martinón-Torres, A. Salas, I. Rivero-Calle, M. Cebey-López, J. Pardo-Seco, J. A. Herberg, N. P. Boeddha, D. S. Klobassa, F. Secka, S. Paulus, R. de Groot, L. J. Schlapbach, G. J. Driessen, S. T. Anderson, M. Emonts, W. Zenz, E. D. Carrol, M. Van der Flier, M. Levin, M. Levin, L. Coin, S. Gormley, S. Hamilton, J. Herberg, B. Hourmat, C. Hoggart, M. Kaforou, V. Sancho-Shimizu, V. Wright, A. Abdulla, P. Agapow, M. Bartlett, E. Bellos, H. Eleftherohorinou, R. Galassini, D. Inwald, M. Mashbat, S. Menikou, S. Mustafa, S. Nadel, R. Rahman, C. Thakker, S. Bokhandi, S. Power, H. Barham, N. Pathan, J. Ridout, D. White, S. Thurston, S. Faust, S. Patel, J. McCorkell, P. Davies, L. Crate, H. Navarra, S. Carter, R. Ramaiah, R. Patel, C. Tuffrey, A. Gribbin, S. McCready, M. Peters, K. Hardy, F. Standing, L. O’Neill, E. Abelake, A. Deep, E. Nsirim, A. Pollard, L. Willis, Z. Young, C. Royad, S. White, P. M. Fortune, P. Hudnott, F. Martinón-Torres, A. Salas Ellacuriaga, F. Álvez González, R. Barral-Arca, M. Cebey-López, M. J. Curras-Tuala, N. García, L. García Vicente, A. Gómez-Carballa, J. Gómez Rial, A. Grela Beiroa, A. Justicia Grande, P. Leboráns Iglesias, A. E. Martínez Santos, N. Martinón-Torres, J. M. Martinón Sánchez, B. Morillo Gutiérrez, B. Mosquera Pérez, P. Obando Pacheco, J. Pardo-Seco, S. Pischedda, I. Rivero-Calle, C. Rodríguez-Tenreiro, L. Redondo-Collazo, S. Serén Fernández, M. del S. Porto Silva, A. Vega, L. Vilanova Trillo, S. B. Reyes, M. C. León León, Á. Navarro Mingorance, X. Gabaldó Barrios, E. Oñate Vergara, A. Concha Torre, A. Vivanco, R. Fernández, F. Giménez Sánchez, M. Sánchez Forte, P. Rojo, J. Ruiz Contreras, A. Palacios, C. Epalza Ibarrondo, E. Fernández Cooke, M. Navarro, C. Álvarez Álvarez, M. J. Lozano, E. Carreras, S. Brió Sanagustín, O. Neth, M. del C. Martínez Padilla, L. M. Prieto Tato, S. Guillén, L. Fernández Silveira, D. Moreno, R. de Groot, A. M. van Furth, M. van der Flier, N. P. Boeddha, G. J. Driessen, M. Emonts, J. A. Hazelzet, T. W. Kuijpers, D. Pajkrt, E. A. Sanders, D. van de Beek, A. van der Ende, R. L. Philipsen, A. O. Adeel, M. A. Breukels, D. M. Brinkman, C. C. de Korte, E. de Vries, W. J. de Waal, R. Dekkers, A. Dings-Lammertink, R. A. Doedens, A. E. Donker, M. Dousma, T. E. Faber, G. P. J. Gerrits, J. A. Gerver, J. Heidema, J. Homan-van der Veen, M. A. Jacobs, N. J. Jansen, P. Kawczynski, K. Klucovska, M. C. Kneyber, Y. Koopman-Keemink, V. J. Langenhorst, J. Leusink, B. F. Loza, I. T. Merth, C. J. Miedema, C. Neeleman, J. G. Noordzij, C. C. Obihara, A. L. T. van Overbeek - van Gils, G. H. Poortman, S. T. Potgieter, J. Potjewijd, P. P. Rosias, T. Sprong, G. W. ten Tussher, B. J. Thio, G. A. Tramper-Stranders, M. van Deuren, H. van der Meer, A. J. van Kuppevelt, A.-M. van Wermeskerken, W. A. Verwijs, T. F. Wolfs, L. J. Schlapbach, P. Agyeman, C. Aebi, C. Berger, E. Giannoni, M. Stocker, K. M. Posfay-Barbe, U. Heininger, S. Bernhard-Stirnemann, A. Niederer-Loher, C. Kahlert, P. Hasters, C. Relly, W. Baer, C. Berger, E. Carrol, S. Paulus, H. Frederick, R. Jennings, J. Johnston, R. Kenwright, C. G. Fink, E. Pinnock, M. Emonts, R. S. Agbeko, S. T. Anderson, F. Secka, K. A. Bojang, I. Sarr, N. Kebbeh, G. Sey, M. Saidykhan, F. Cole, G. Thomas, M. Antonio, W. Zenz, D. S. Klobassa, A. Binder, N. A. Schweintzger, M. Sagmeister, H. Baumgart, M. Baumgartner, U. Behrends, A. Biebl, R. Birnbacher, J.-G. Blanke, C. Boelke, K. Breuling, J. Brunner, M. Buller, P. Dahlem, B. Dietrich, E. Eber, J. Elias, J. Emhofer, R. Etschmaier, S. Farr, Y. Girtler, I. Grigorow, K. Heimann, U. Ihm, Z. Jaros, H. Kalhoff, W. Kaulfersch, C. Kemen, N. Klocker, B. Köster, B. Kohlmaier, E. Komini, L. Kramer, A. Neubert, D. Ortner, L. Pescollderungg, K. Pfurtscheller, K. Reiter, G. Ristic, S. Rödl, A. Sellner, A. Sonnleitner, M. Sperl, W. Stelzl, H. Till, A. Trobisch, A. Vierzig, U. Vogel, C. Weingarten, S. Welke, A. Wimmer, U. Wintergerst, D. Wüller, A. Zaunschirm, I. Ziuraite, V. Žukovskaja, Life-threatening infections in children in Europe (the EUCLIDS Project): a prospective cohort study. The Lancet Child & Adolescent Health 2, 404–414 (2018).

47. P. Bankhead, M. B. Loughrey, J. A. Fernández, Y. Dombrowski, D. G. McArt, P. D. Dunne, S. McQuaid, R. T. Gray, L. J. Murray, H. G. Coleman, J. A. James, M. Salto-Tellez, P. W. Hamilton, QuPath: Open source software for digital pathology image analysis. Sci Rep 7, 16878 (2017).

48. A. E. Carpenter, T. R. Jones, M. R. Lamprecht, C. Clarke, I. H. Kang, O. Friman, D. A. Guertin, J. H. Chang, R. A. Lindquist, J. Moffat, P. Golland, D. M. Sabatini, CellProfiler: image analysis software for identifying and quantifying cell phenotypes. Genome Biol 7, R100 (2006).

