## Supplementary Material for "Bacterial histone-like proteins released during antibiotic treatment mediate vascular injury in meningococcal sepsis"

**Affiliations**:

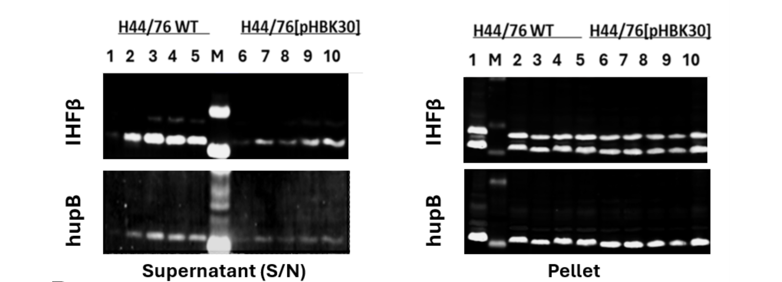
**A**

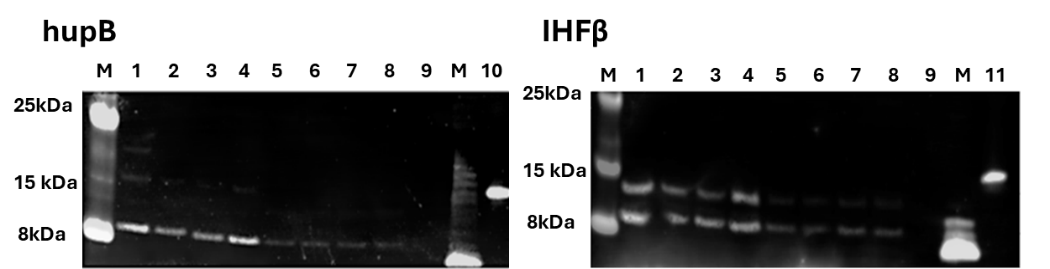

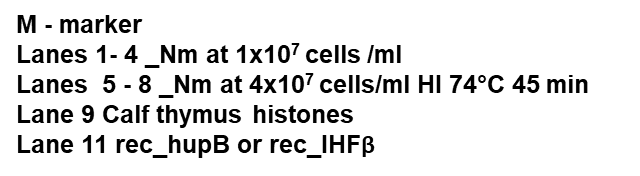

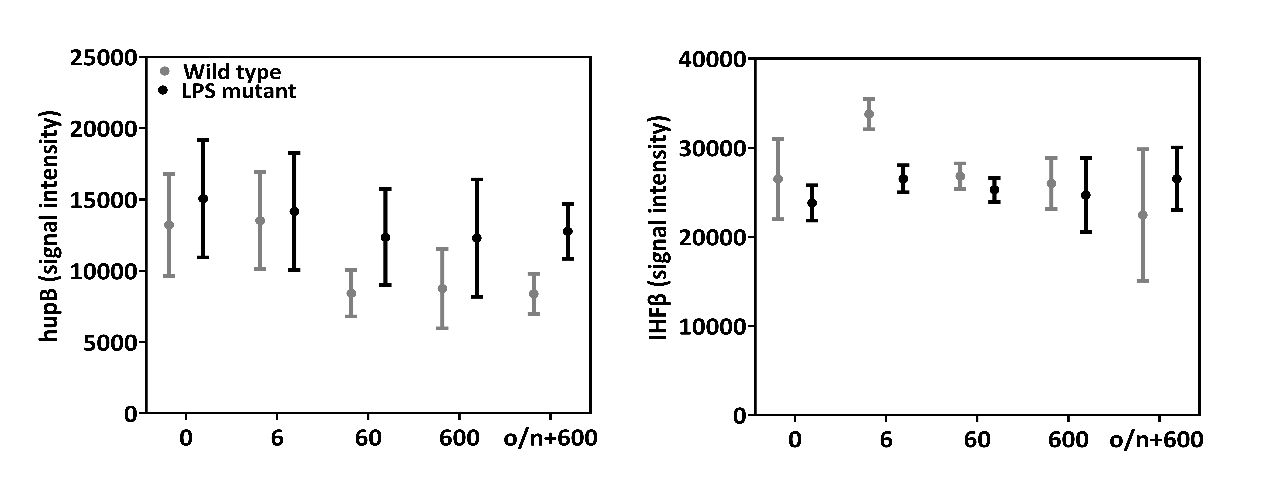

**B**

**figure S1. (A)** Histone like proteins (HLPs) are detectable in supernatants and pellets from *Neisseria meningitidis* cultures after treatment with Ceftriaxone using Western blot. Lines 1,6 – no Ceftriaxone; lines 2,7 – 6 ng/ml; lines 3,8 – 60 ng/ml; lines 4,9 – 600 ng/ml and lines 5,10 – overnight 600ng/ml); Antibody specificity was also confirmed using *Neisseria meningitidis* M58 strain **(B)** The levels of HLPs in the *Neisseria meningitidis* pellets are not dependent on Ceftriaxone doses. Data presented as mean ± SD.

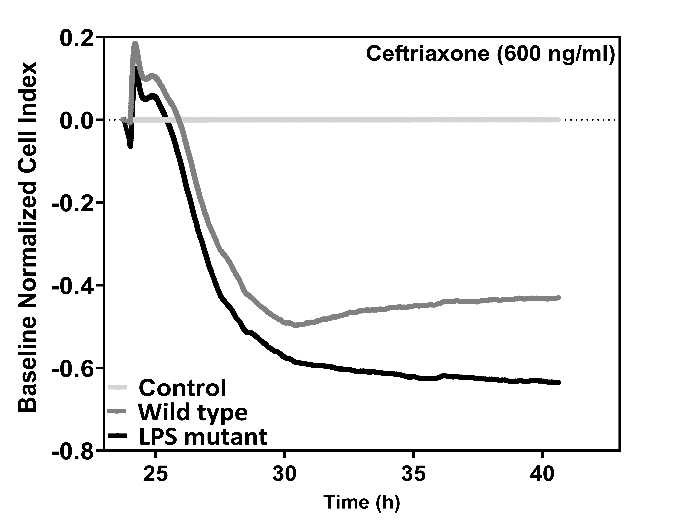

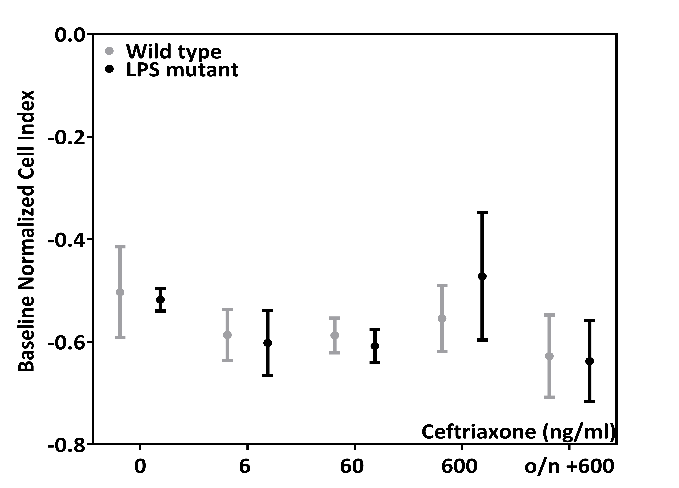

**figure S2.** The pellets from *Neisseria meningitidis* cultures after treatment with Ceftriaxone affect human primary dermal endothelial cells (HDMVEC) permeability, however this effect is not dose related.

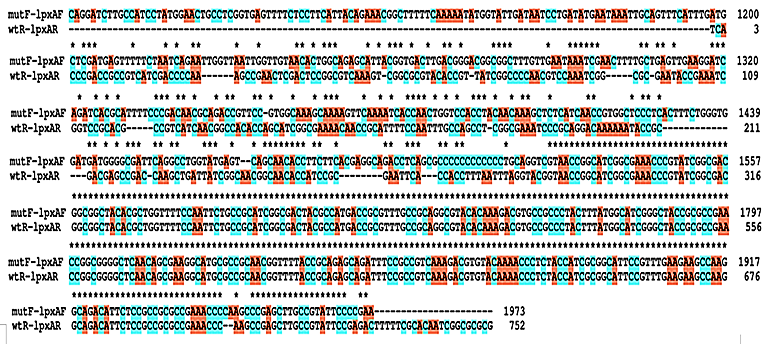
**(A)**

**(B)**

| **Rapid gel clot Endotoxin Assay (0.125 U/ml)** | | | |
| --- | --- | --- | --- |
| **Sample ID** | **PC** | **Result** |  |
| Media +Ceftriaxone | + | - | Negative |
| Wild type | + | + | Endotoxin level >0.125 U/ml |
| Wild type + Ceftriaxone | + | + | Endotoxin level >0.125 U/ml |
| LPS mutant | + | - | Negative |
| LPS mutant + Ceftriaxone | + | - | Negative |

**figure S3. (A)** Comparison of LPS locus between used bacterial strains. **(B)** Detection of Endotoxin levels in the *Neisseria meningitidis* culture components. PC -positive control.

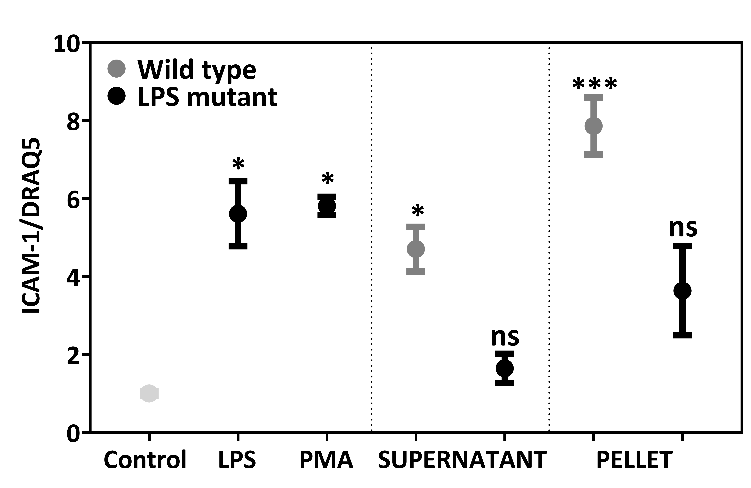

**figure S4.** *Neisseria meningitidis* culture components activate endothelial cells. LPS-EB (LPS from *E. coli* O111:B4; 75 ng/ml); PMA (Phorbol 12-myristate 13-acetate, 5nM). *P < 0.05, **P < 0.01; ****P < 0.0001 statistically significant. Experiments were performed in triplicate. Data presented as Mean with SD.

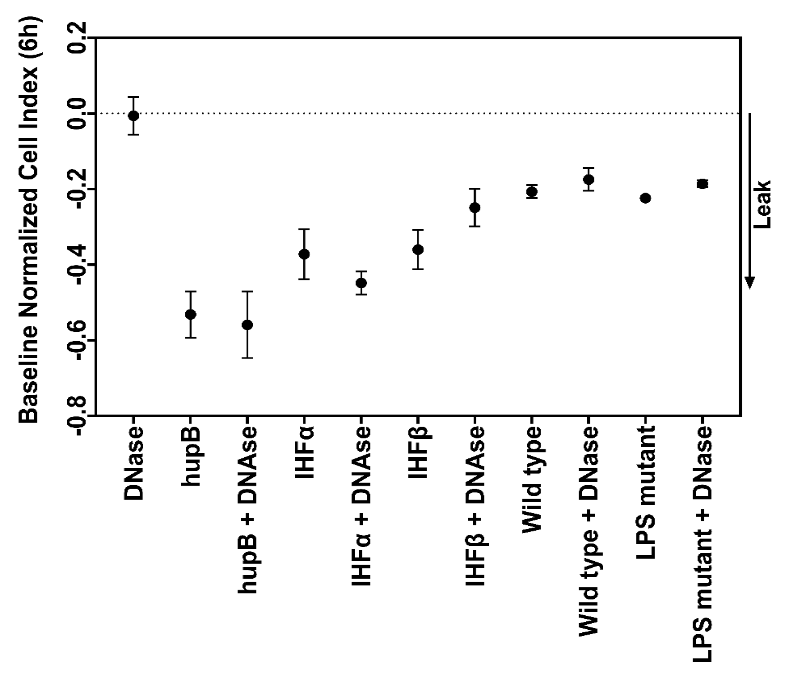

**figure S5.** DNase I (10 U/ml of 25μg/ml protein sample) treatment doesn’t have an effect on recombinant histone-like proteins (HLPs) mediated HDECs permeability. DNA removal was performed using RQ1 RNase-Free DNase kit (Promega), followed by cleanup of samples using ultrafiltration columns (10 kDa MWCO) and DNA content confirmation (Qubit dsDNA HS kit; ThermoFisher)

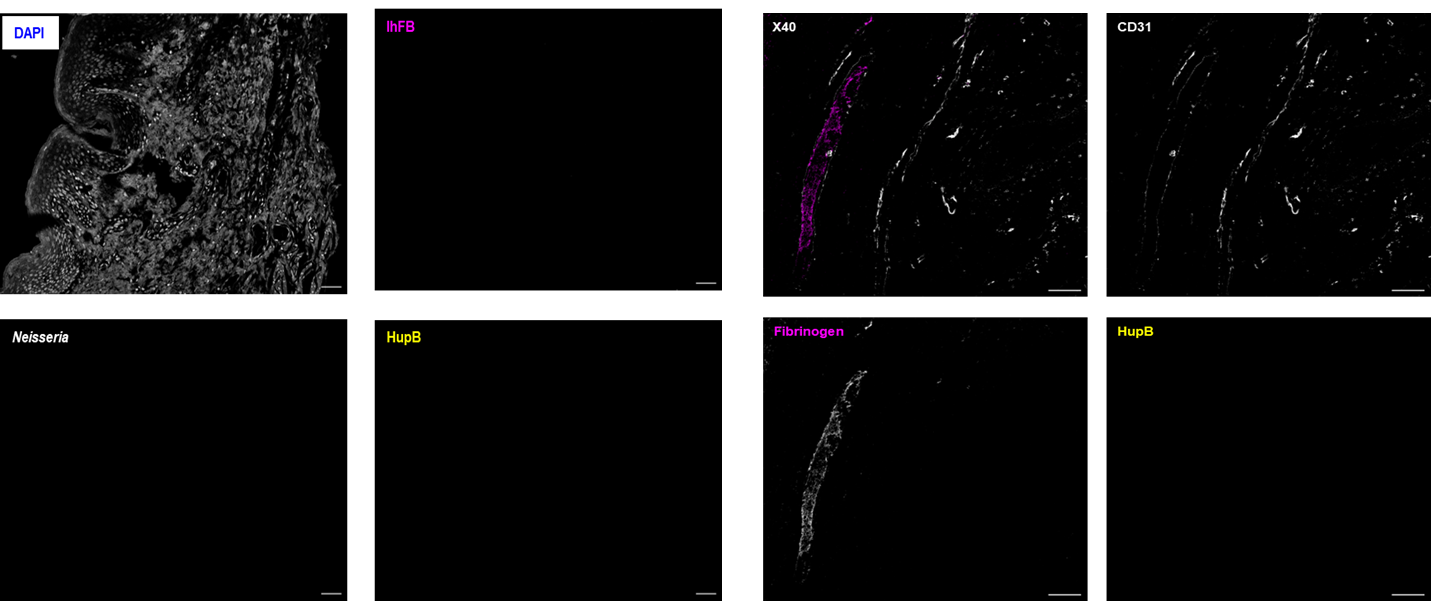
**(A)	(B)**

**figure S6. (A)** Examples of staining for *Neisseria meningitidis* and HLPs in control skin punch biopsies. **(B)** Examples of staining for Fibrinogen and HLPs (hupB; IHFβ) in control skin punch biopsies. CD31- marker of endothelial cells

**(B)**

**(A)**

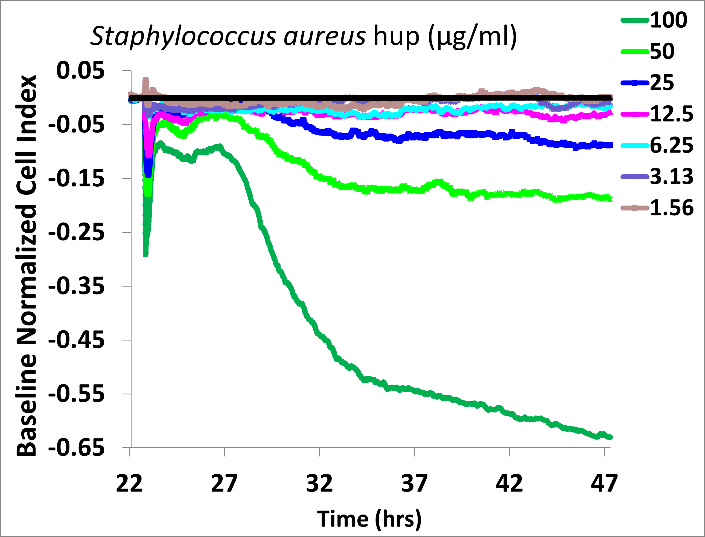

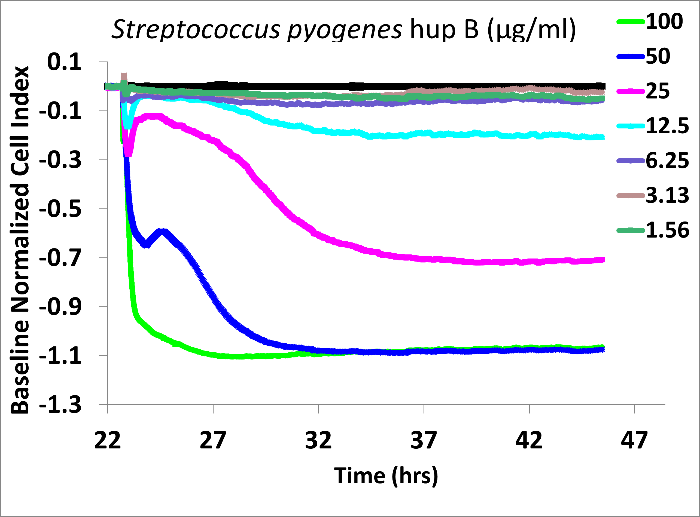

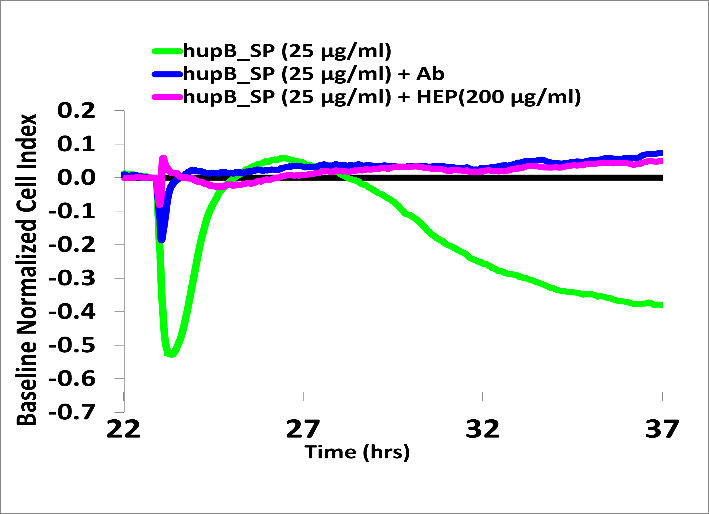

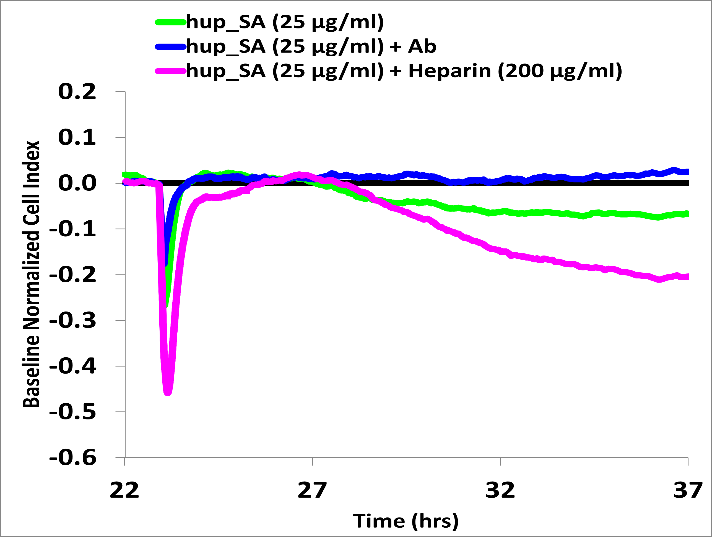

**figure S7**. Endothelial cells permeability is affected by hup from Staphylococcus aureus (SA) and hupB from Streptococcus pyogenes (SP). **(A)** Serial dilutions of recombinant proteins **(B)** The effect of pre-incubation of HLPs with non-anticoagulant N-Acetylheparin (HEP, 200 μg/ml). Anti His-tag antibodies (250 ng) were used as positive controls to remove recombinant HLPs.

**figure S8.** The effect of recombinant *Neisseria meningitis* HLPs (HupB, IHFα and IHF β), Hu proteins from Staphylococcus aureus (SA, hup) and Streptococcus pyogenes (SP, hupB) on *Galleria mellonella* larvae survival is dose dependent. Calf thymus histones (CTH) were used as a positive control. Each experimental group consisted of 10 larvae without signs of melanisation; experiments were performed with 3 biological replicates of supernatants and controls on at least 3 occasions. Asterisks indicate significant differences relative to the control group of waxworm and treatment groups, as assessed by the log-rank (Mantel–Cox) test (ns, non-significant, *P < 0.05, **P < 0.01; ****P < 0.0001).

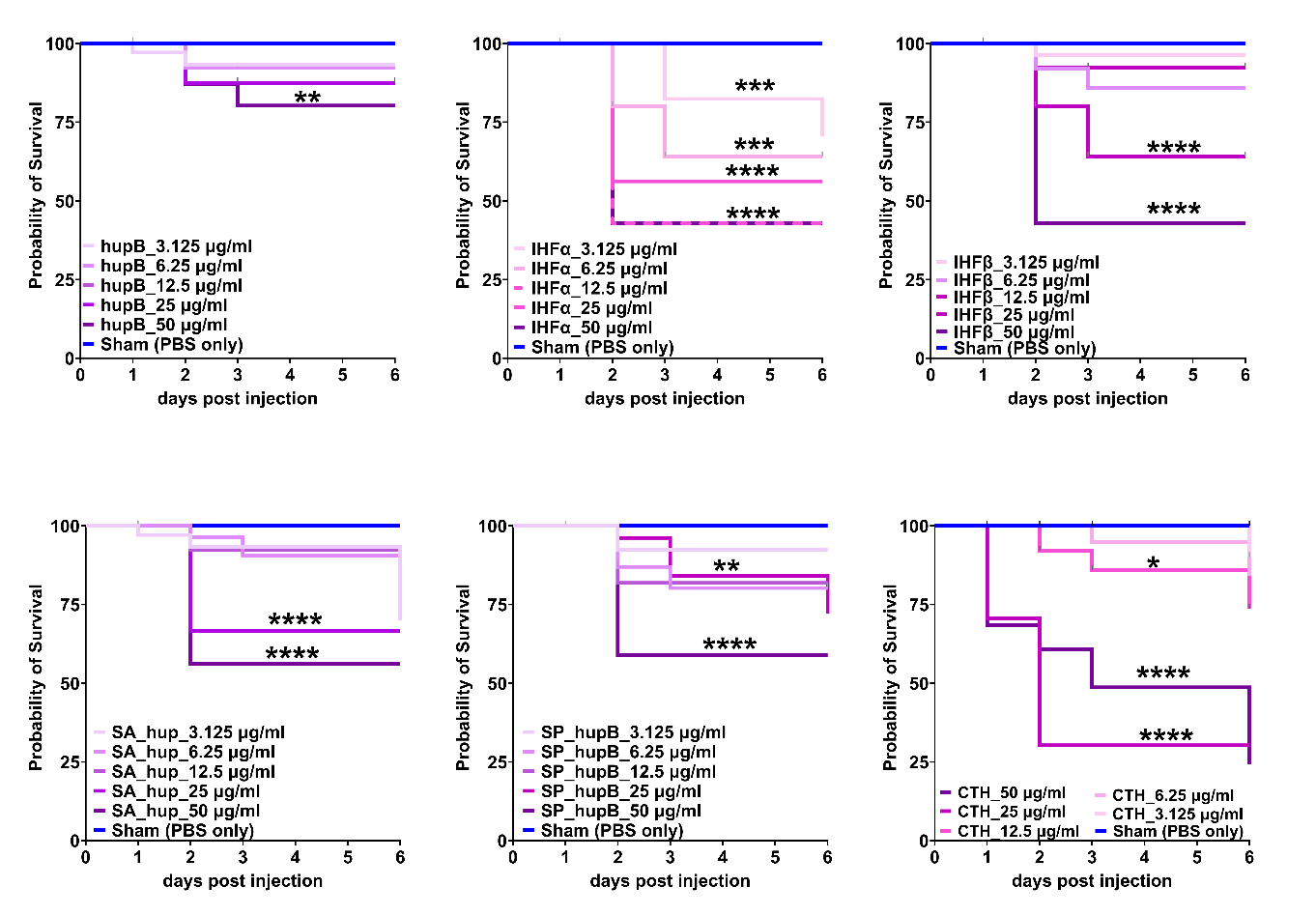

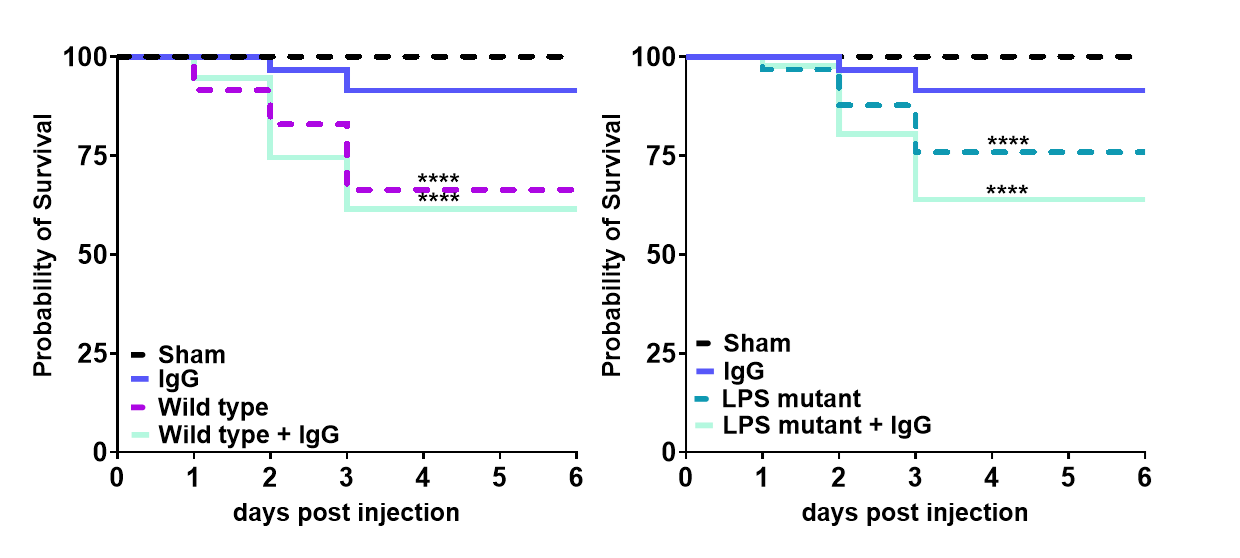

**figure S9.** Presence of rabbit IgG (250 ng) did not reduce supernatant toxicity or improved *Galleria mellonella* larvae survival. Asterisks indicate significant differences relative to the IgG treated waxworm and treatment groups, as assessed by the log-rank (Mantel–Cox) test (ns, non-significant, *P < 0.05, **P < 0.01; ****P < 0.0001).

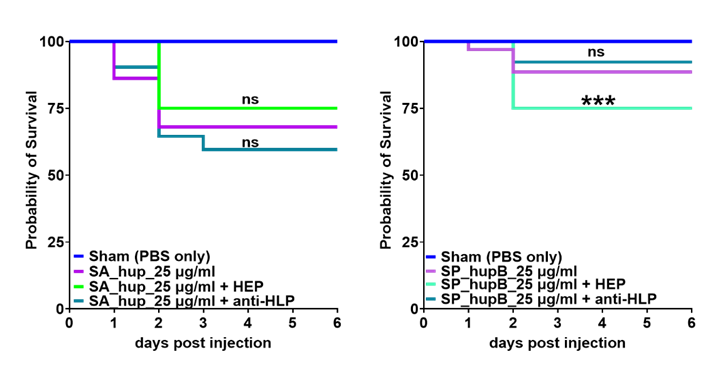
**figure S10.** The effect of non-anticoagulant N-Acetylheparin (HEP, 200 μg/ml)) or cocktail of the anti-HLP antibodies (250 ng) the on *Galleria mellonella* larvae survival after the treatment with the recombinant Hu proteins from Staphylococcus aureus (SA) and Streptococcus pyogenes (SP). Each experimental group consisted of 10 larvae without signs of melanisation; experiments were performed with 3 biological replicates of supernatants and controls on at least 3 occasions. Asterisks indicate significant differences relative to the HLPs, as assessed by the log-rank (Mantel–Cox) test (ns, non-significant, *P < 0.05, **P < 0.01; ****P < 0.0001).

**Supplementary methods (SM)**

**Trans endothelial electrical resistance (TEER) analysis**

Endothelial barrier function was measured using xCELLigence RTCA S16 or DP (Agilent, US), were seeded at 40,000 cells/well in 16 well PET E-plates (surface area 0.2 cm^2^) coated with the bovine fibronectin (10μg/ml, ThermoFisher, UK; 33010018), transendothelial electrical resistance (TEER) was recorded and expressed as cell index. The decrease in the endothelial barrier function was determined by normalising the cell index at the point immediately before adding treatment and further normalised to the baseline (control sample) giving read out as the baseline normalised cell index (BNCI). Briefly, HDMECs were grown overnight until confluency was established (cell index 10 or above) in the EGM-MV with supplements, then media was exchanged for OptiMEM^TM^ medium (ThermoFisher; 31985062) supplemented with 1% KnockOut™ Serum Replacement (ThermoFisher; 10828010) for 2 h before treatments alone or in combinations were added **(table SM1)**.

**Measurement of cell viability**

Endothelial cell viability was measured using Apotracker™Green (Apo-15 peptide) a calcium-independent probe for detecting apoptotic cells (BioLegend; 427403) in the combination with the Live-or-Dye™594/614 (Biotium; 32006) and Hoechst 33342 (ThermoFisher; 62249). Briefly, HDMECs cells were seeded at 64,000 cells/well (surface area 0.32 cm^2^) in the attachment factor coated 96 well optical plates and grown until reaching confluency in the EGM-MV media, then exchanged for OptiMEM^TM^ supplemented with 1% KnockOut™Serum Replacement for 2h before treatments. Positive controls were created by treatment of cells with PMA (5 nM), TNFα (100 ng/ml) or Staurosporine (1 μM). Both the Apotracker™Green and Live-or-Dye™594/614 stains were diluted in the Opti-MEM^TM^ medium supplemented with 1% KnockOut™ Serum to 1:500 dilution and 100 μl of that solution was added to 100 μl of media already present in the plate giving final dilution of 1:1000. Cells were incubated protected from the light for 30 min, during the last 5 min Hoechst 33342 was added (2 μg/ml) to visualise nuclei. After incubation cells were washed in Ca^2+^Mg^2+^free PBS, fixed in 4% methanol-free paraformaldehyde in Ca^2+^Mg^2+^free PBS for 10 min in RT, washed with PBS and imaged immediately on the 96 well plate. Each stain was performed in duplicate with 5 fields being used for analysis each time. QuPath software was used to evaluate endothelial cells viability.

**table SM1. List of compounds and working concentrations used in the study.**

| **Compound** | **Compound name** | **Supplier** | **Concentration used** |
| --- | --- | --- | --- |
| PMA | Phorbol 12-myristate 13-acetate | InvivoGen, tlrl-pma | 5nM |
| LPS | LPS-EB (LPS from E. coli O111:B4) | InvivoGen; tlrl-eblps | 75 ng/ml |
| TNFα | Recombinant Human TNF-alpha (HEK293-expressed) Protein, CF | R&D Systems; 10291 | 100 ng/ml |
| HKB | Heat Killed E. coli 0111:B4 | InvivoGen; tlrl-hkeb2 | 10^8^ cells/ml |
| HupB | Recombinant Neisseria meningitidis serogroup B DNA-binding protein Hu-beta (hupB) | Cusabio;  CSB-EP350724NGG | 25 μg/ml |
| IHFα | Recombinant Neisseria meningitidis serogroup B Integration host factor subunit alpha (ihfA) | Cusabio;  CSB-EP351432NGG | 25 μg/ml |
| IHFβ | Recombinant Neisseria meningitidis serogroup B Integration host factor subunit beta (ihfB) | Cusabio;  CSB-EP357958NGG | 25 μg/ml |
| Heparin/HEP | N-Acetylheparin sodium salt | Merck; A8036 | 200 μg/ml |
| Hp-A | Hp-A | Intellihep Ltd | 200 μg/ml |
| Hp-B | Hp-B | Intellihep Ltd | 200 μg/ml |
| Hp-C | Hp-C | Intellihep Ltd | 200 μg/ml |
| Hp-C | Hp-C | Intellihep Ltd | 200 μg/ml |
| HS | Heparan sulphate | MedChemexpress LLC; HY-101916 | 200 μg/ml |

**Immunofluorescence**

Briefly, HDMECs cells were seeded at 200,000 cells/cm^2^ in the attachment factor coated culture dishes (96 well optical plates or 8 well chambers) or and grown until reaching confluency in the EGM-MV media, then media was exchanged for Opti-MEM^TM^ with 1% KnockOut™Serum Replacement for 2h before treatments. After treatment cells were fixed immediately in 4% methanol-free paraformaldehyde in Ca^2+^Mg^2+^free PBS for 10 min in RT, washed with 3 times with PBS, incubated with 100mM Glycine/PBS for 7 min RT, washed 3x in PBS. When required for intracellular staining, cells were permeabilised with 0.1% Triton X100 in PBS for 10 min in RT, washed 3x in PBS and incubated with the Intercept®(PBS) Blocking Buffer (Li-COR, P/N: 927-70001) with addition of 5% goat serum for 1h in RT, primary antibodies against HLPs were diluted 1:1000 in Intercept®T20 (PBS) Antibody Diluent (Li-COR; P/N: 927-75001) and incubated o/n at 4°C. After incubation plates/chambers were washed 3x with PBST (PBS-0.1%Tween 20), followed by an 1h incubation with Goat anti-Rabbit IgG (H+L), Alexa Fluor™ Plus 555 (1:2000, ThermoFisher UK, # A32732), and 3 x PBST washes. Then cells were stained with endothelial markers namely cells were stain with anti-human CD201/EPCR (1:100; BioLegend, cat no 351905), anti-human CD144/VE-cadherin (1:250; Miltenyi Biotec, cat no: 130-118-358), anti-human CD31 (1:200; Miltenyi Biotec, cat no. 130-110-668) or CD54 (1:100 Miltenyi Biotec, cat no. 130-120-711). If different conjugated fluorophores were used for a particular staining, the same antibody dilution was applied for IF staining. Cells were counterstain with Hoechst 33342. Images were acquired on EVOS FL2 microscope directly from the chambers/wells and saved as grey scale pictures. QuPath software was used to evaluate endothelial cells surface markers.

**Western blot**

Plasma samples (3ul) were mixed with NuPAGE^TM^LDS sample buffer in the presence of NuPAGE^TM^Reducing Agent and heated at 70°C for 10 minutes. Proteins were then separated on 12% Bis-Tris gels in MES-SDS running buffer and transferred onto Immobilon-FL PVDF membranes (LI-CORE, UK). Membranes were blocked in Intercept® Blocking Buffer PBS for 1h RT (LI-CORE, UK), followed by the incubation o/n at 4°C with custom primary antibodies: rabbit-anti *Neisseria meningitidis* serogroup B DNA-binding protein hu-beta (hupB; 1:5000; Cusabio), or rabbit-anti IHFα (1:5000; Cusabio) or rabbit-anti IHFβ (1:5000; Cusabio) diluted in Intercept PBS T20 Antibody Diluent (LI-CORE, UK), membranes were washed with PBS-T (0.2% Tween 20) and incubated for 1h RT with secondary antibodies, IRDye®800CW or IRDye®680RD donkey anti-rabbit IgG (1:20,000 dilution in PBS/0.2%Tween 20/ 0.01% SDS, LI-CORE, UK). Membranes were washed PBS-T (0.2% Tween 20), followed by washes in PBS to remove detergent before visualisation using Odyssey CLx imaging system and Image Studio software (LI-CORE, UK).

**In-Cell Western**

Briefly, HDMECs cells were seeded at 40,000 cells/well in the attachment factor coated 96 well plates for In-Cell Western (Li-COR, UK; P/N: 926-19156) and grown until reaching confluency in the Endothelial Cell Growth Medium MV with supplements, then media was exchanged for OptiMEM^TM^ medium supplemented with 1% KnockOut™ Serum Replacement for 2h before treatments. After treatment cells were fixed immediately in 4% methanol-free paraformaldehyde for 10 min in RT, washed with PBS then permeabilised with ice-cold methanol for 10 min and washed again in 3x PBS. Plates were blocked using Intercept®(PBS) Blocking Buffer (Li-COR, P/N: 927-70001) for 1h in RT, primary antibodies were diluted in Intercept®T20 (PBS) Antibody Diluent (Li-COR; P/N: 927-75001) and incubated o/n at 4^°^C. The final concentration of antibodies used were CD144/VE-cadherin 1:1000 (Abcam; ab33168), CD201/EPCR 1:500 (1489; Emson) and CD54/ICAM-1 (1:250). After incubation plates were washed 3x with PBST (PBS-0.1%Tween 20). IRDye800CW conjugated secondary goat anti-rabbit (Li-COR; P/N: 926-32211) or goat anti-mouse (Li-COR; P/N: 926-32210) antibodies were used at 1:1000 dilution and incubated for 1h in RT in the dark. For cell number normalisation DRAQ5™ (Biostatus Ltd; DR05500, 1:10,000) and Sapphire700™stain (Li-COR; P/N: 928-40022 at 1:1000) dyes were mixed and added together with secondary antibody mixture. Background wells were created by omitting primary antibodies and cell stains as additional control unstained cells were also added. After incubation plates were washed 3x in PBST and filled with 100 μl of PBS. Plates were scanned on an Odyssey®CLx Imaging System (Resolution: 169 µm; Focus offset: 3.5 mm; Intensity: Auto for both channels). Data were analysed using Empiria Studio Software and presented as a ration of ICW of target normalised to cell number.

**Immunohistochemistry**

FFPE skin samples from children with meningococcal disease and controls were stained using a multiplex Tyramide Signal Amplification system (Opal-TSA, Akoya Biosciences). 3μm sections were dewaxed and rehydrated sequentially in Xylene (2x 7 min) followed by series of washes 2x 2 minutes in each of ethanol solutions (absolute alcohol, 95 %, 90 % and 70 % ), rinsed in water for 5–10 min. Heat-induced antigen retrieval performed **(table SM2),** endogenous peroxidase/pseudoperoxidase activities inhibition (BLOXALL, Vector laboratories) were performed sequentially. Sections were incubated with Fc Receptor Blocking Solution (Human TruStain FcX™, BioLegend) followed by a blocking step for 1h RT (5% human serum, 5% BSA and 10% serum corresponding to the host of secondary antibody). Incubation with primary antibodies diluted in blocking solution was performed overnight at (4°C). Detailed information about primary antibodies and concentrations used in this study are described in **table SM2**. Primary antibodies were incubated with species-specific HRP conjugated secondary antibodies (ImmPRESS®HRP IgG Polymer Detection kit, Vector Laboratories) and were visualised with Opal-TSA, see **table SM2**). When multiplexing, heat induced antigen retrieval was performed to strip primary/secondary antibody pairs, while preserving the antigen-associated fluorescence signal. Tissue specific optimisation for all antibodies was conducted on control tissue. Isotype controls were used to ensure specificity.

**table SM2.** Antibody information and working concentration, antigen retrieval methods and other information regarding IHC.

| **Antibody** | **Supplier (cat no)** | **Antigen retrieval** | **Antibody concentration** | **Notes** |
| --- | --- | --- | --- | --- |
| CD31 (intracellular) | Novus  (NB100-2284) | Cit pH6  (90min@60c oven) | 2 μg/mL | TSA time 10 min  FCR block 5 min |
| Fibrinogen | Invitrogen  (PA1-26809) | EDTA pH9  Pressure Cooker | 6 μg/mL | TSA time 5 min  FCR block 5 min |
| HupB | Cusabio (CSB-A350724XA01NGG) | Cit pH6  (90min@60c oven) | 2 μg/mL | TSA diluted 1:200, FCR block 10 min |
| IhFA | Cusabio (CSB-A351432XA01NGG) | Cit pH6 (90min@60c oven) | 1 μg/mL | TSA diluted 1:200, FCR block 10 min |
| IhFB | Cusabio (CSB-A357958XA01NGG) | Cit pH6 (90min@60c oven) | 2 μg/mL | TSA diluted 1:200, FCR block 10 min |
| Neisseria meningitidis | LSBio  (LS‑C683288) | Cit pH6  Pressure Cooker | 2 μg/mL | TSA diluted 1:200, FCR block 10 min |
| EPCR | R&D Systems  (AF2245) | EDTA pH9 (MWT-Boil-10min@10%) | 0.02 μg/mL | TSA time 10 min  FcR block 20mins |

**Correlative light-electron microscopy (CLEM)**

CLEM approach was used to investigate co-localisation of HLPs and Neisseria bacteria in the meningococcal tissue biopsies. Briefly, coverslips were later removed by floating them away through addition of water. Tissue sections were re-fix in 2.5% glutaraldehyde and 4% paraformaldehyde in 0.1M cocodylate buffer. After 3 washes in the same buffer, the tissue sections were dehydrated in ascending ethanol series (30%, 50%, 70% and 100%) for 10 minutes in each step. The sections were then dried using HMDS – 4 x 5 minutes each. The dried samples were mounted onto stubs by fixing the slides using a carbon tape. Slides were covered with 20nm layer of gold/palladium using a Quorum Q1-ES metal coater.
